# Human-specific multicopy gene FRMPD2 promotes synapse formation via recruitment of neuroligin 1

**DOI:** 10.64898/2026.04.07.716942

**Authors:** Yuda Huo, Ameya Patkar, KathrynAnn Merth, James Gilbert, Heng-Ye Man

## Abstract

FRMPD2 is a human-specific multi-copy gene with higher mRNA expression in brain tissue, but its role in synapse formation and neurodevelopment remains unknown. We find that FRMPD2 is a neuron-specific protein with a high expression level in human brains compared to rodent brains. FRMPD2 overexpression in rat neurons stimulates synaptogenesis, leading to increased synaptic activities. Importantly, our results show that FRMPD2 via its PDZ domains recruits and enhances neuroligin-1 protein levels at the postsynaptic sites, and via its FERM domain interacts with the F-actin network in the spine. Increased expression of FRMPD2 also promotes spine formation and maturation, a foundational process for synapse formation. In developing embryonic mouse brains expressing higher FRMPD2 protein levels, we observed delayed neuronal migration, presumably promoting a protracted timeline for cortical lamination as a feature in human brain development. Behaviorally, mice with FRMPD2 overexpression in the brain demonstrate enhanced spatial memory retention. These findings indicate an important function for FRMPD2 in neuronal connectivity, brain development, and cognitive function.

## INTRODUCTION

Synaptogenesis is a highly regulated developmental process that can be divided into multiple steps, including initial synaptic contact, target recognition, stabilization, and functional maturation via recruitment and assembly of the pre- and postsynaptic molecular machinery (Südhof 2018, Sheng 2011). Cellular adhesion molecules (CAM) play a key role in cell-cell connection and closely participate in synapse formation and assembly of pre- and postsynaptic proteins (Brose 1999, Missler 2012, Biederer 2002, Sheng 2011, Südhof 2011). Synaptically localized CAMs have been shown to facilitate this process by direct recruitment of intracellular functional and structural elements through their intracellular terminals, particularly PDZ binding domains (Mondin 2011, Dalva 2007, Waites 2005). Of the identified synaptic CAMs, neurexin and neuroligin, which are distributed at the pre- and the postsynaptic site, respectively (Nguyen 1997, Südhof 2008), act as central organizers during synapse formation (Graf 2004, Nam 2005, Chih 2005, Poulopoulos 2009). They not only mediate specific synaptic contacts but also directly recruit and assemble key synaptic components, including synaptic vesicles, neurotransmitter receptors, and postsynaptic scaffolds (Irie 1997, Craig 2007, Bukalo 2012, Shipman 2012, Futai 2013). Heterologous synaptogenic models have been established to study the effects of various protein complexes on synapse formation in non-neuronal cells. CAMs like neuroligin, neurexin, and synCAM play an important role during initiation of synaptogenesis through recruitment and assembly of synaptic elements (Scheiffele 2000, Biederer 2002, Yamagata 2018). In co-cultures of both neurons and HEK cells, expression of neuroligin in HEK cells is sufficient to attract axon terminals and induce presynaptic maturation and synaptic vesicle accumulation (Scheiffele 2000, Biederer 2002), whereas expression of neurexin in cells results in recruitment of glutamate receptors and postsynaptic assembly (Graf 2004, Nam 2004).

The family of FERM and PDZ-domain containing (FRMPD) proteins has seven members, which belong to the FERM protein superfamily (An 2008, Piard 2018, Wang 2020). FRMPD proteins have been studied with respect to synaptic plasticity in response to activity and remodeling through downstream signaling. Specifically, FRMPD1 is implicated in adaptive translocation of the G-protein transducin, thereby inducing activity-dependent GPCR signaling to modulate rod photoreceptor synapses (Campla 2019, Campla 2022). FRMPD1 is also shown to regulate the subcellular localization of AGS3, a protein involved in neural development and synaptic plasticity (Ningfei 2008). FRMPD3, another member of this family, is implicated in regulating susceptibility to epilepsy via enhancing surface integration of the GluA2 subunit of AMPA receptors (AMPARs) and forming a complex with GRIP. Knockdown of FRMPD3 reduces neuronal excitability and epileptogenic cellular physiology, indicating the function of the FERM family proteins as regulatory proteins at excitatory synapses (Jia 2025). Genetic studies have also indicated FRMPD3 as a potential risk gene for ASD and MDD (Ben-Mahmoud 2024, Oh 2024). While the specific mechanisms remain unresolved, FRMPD3’s interaction with Npas3 in regulating the excitation/inhibition balance and activity-dependent cell signaling further emphasizes the regulatory functions of FRMPD proteins at neuronal synapses. Deleterious mutations in FRMPD4 are implicated in X-linked intellectual disability (ID) and a cellular phenotype of impaired dendritic spine morphogenesis (Pan 2024, Li 2024, Piard 2018, Lee 2008). FRMPD4 has been reported to interact with metabotropic glutamate receptors through the FERM domain (Wang 2020), and intronic variants lead to exon skipping the FERM domain, leading to a patient with ID (Satake 2025). These findings suggest an important role for FRMPD proteins in brain development and synaptic functions (Sudmant 2010, Dennis 2017).

FRMPD2 (FERM and PDZ domain containing protein 2) is initially predicted to be a novel scaffolding protein from the genome (Deloukas 2004, Gerhard 2004, Ota 2004). It consists of one kinase non-catalytic C-lobe domain (KIND) on the N-terminal, followed by 4.1N/Ezrin/Radixin/moesin (FERM) domain, and three PDZ domains at the C-terminal. While all of the 7 members in the FRMPD family share the FERM domain (with no other domains in FRMPD3-7), only FRMPD1/2 are PDZ-containing, with one PDZ domain in FRMPD1 and three PDZs in FRMPD2. The specific biological roles and molecular mechanisms of FRMPD2 in the brain and neurons remain largely unknown. Studies in epithelial cells indicate that FRMPD2 is localized in a polarized manner to tight junctions through PDZ domain-mediated association with E-cadherin (Stenzel 2009), suggesting a role in intercellular connection and communication (Seong 2001). In neurons, PDZ domain-containing proteins often function as scaffolding molecules at the synaptic site. A structural feature for post-synaptic scaffold proteins is the presence of a single or multiple PDZ domains (Hata 1998, Kim 2004, Buday 2010, Erlendsson 2019). Scaffolding proteins can thus associate with other proteins containing a PDZ-binding motif and enable assembly of the scaffold network in the postsynaptic compartment (Liu 2019, Beuming 2005). Given that FRMPD2 contains three PDZ domains, it is predicted to be an ideal molecule to form multiple interactions (Harris 2001, Lu 2019) and potentially play a role in synaptic organization. Evolutionarily, FRMPD2 has been identified as one of the human-specific multi-copy genes with 2 copies in the human genome (Sudmant 2010). Both the ancestor and the duplicated FRMPD2 gene copies show temporally controlled mRNA expression in both developing and adult human brains (Dougherty 2018, Soto 2025). The functional significance of high-dosage expression of FRMPD2 during human brain development remains unclear.

We report that the FRMPD2 protein is expressed preferentially in the brain, with a higher expression in human brains compared to mouse brains. We found that FRMPD2 is selectively localized at excitatory synapses, and overexpression of FRMPD2 promotes synaptogenesis. Mechanistically, FRMPD2 is targeted to synapses through binding of its FERM domain to F-actin within dendritic spines. Through its PDZ domains, FRMPD2 further interacts with the synaptic adhesion molecule neuroligin-1, enhancing its surface expression and consequently facilitating synaptogenesis. Overexpression of the FRMPD2 protein in mouse brains results in an increase in dendritic spine density, accompanied by a delay in neuronal migration. Importantly, FRMPD2 overexpression in the brain leads to stronger spatial memory retention in mice.

## RESULTS

### FRMPD2 is specifically expressed in neurons and shows elevated protein level in human brains

Previous genetic studies indicate that the FRMPD2 gene is duplicated on human chromosome 10 and the duplication occurs only in the human genome but not in other primates, resulting in two copies in the human genome (Sudmant 2010, Dennis 2017). To examine the expression of FRMPD2 protein levels, we performed western blotting of cortical brain lysates from human brains and age-matched rat brains. Interestingly, we found that the FRMPD2 protein level in human brains is 1-fold higher than that in rat brains (**Fig. 1A-B**). PSD-95 was probed as a negative control and showed no change between humans and rats.

**Figure 1.**
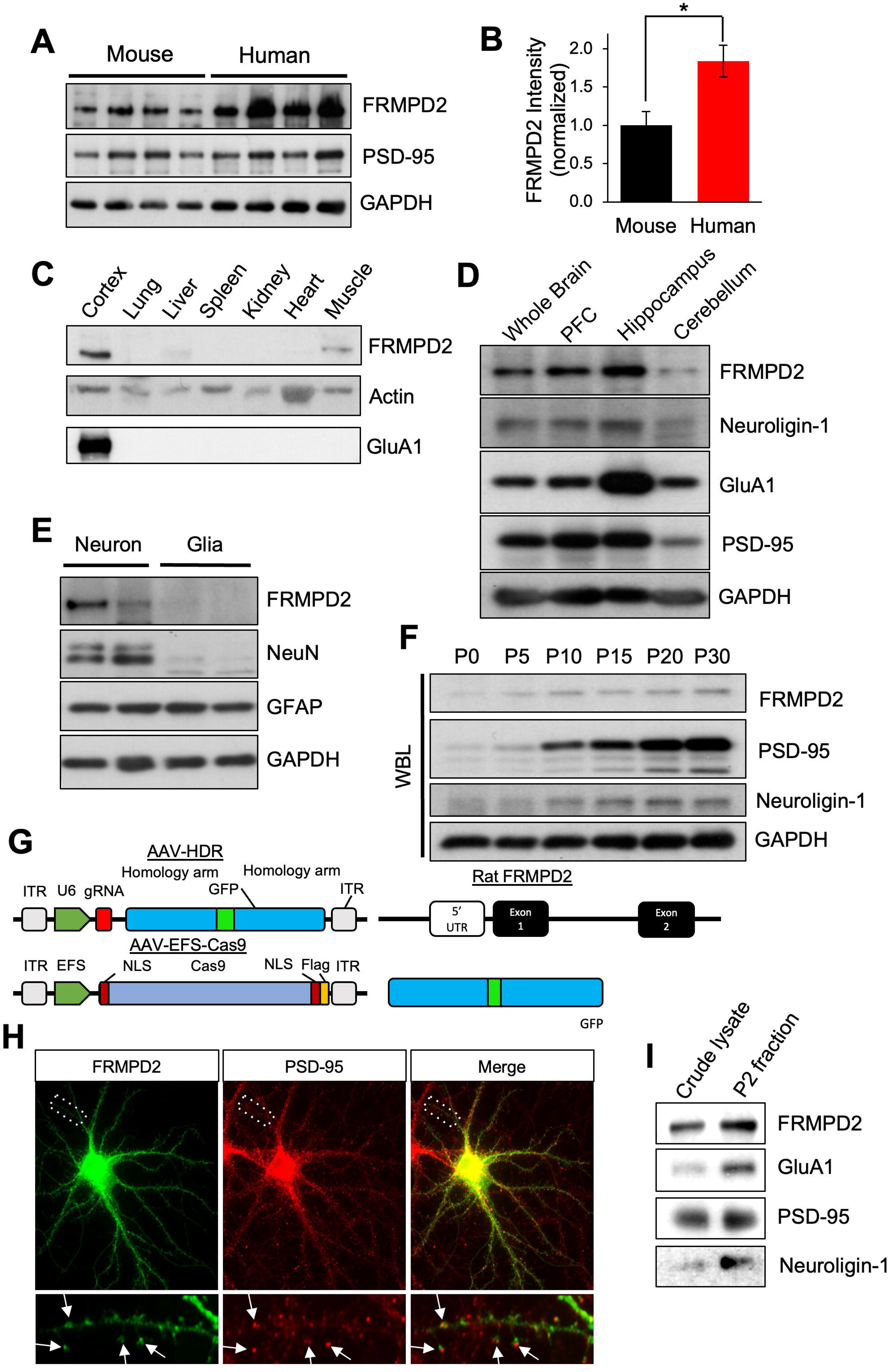
FRMPD2 protein expression profiling and characterization of *in vitro* localization. **(A)** Protein expression profiling *in vivo* shows that FRMPD2 is a neuron-specific protein with higher expression levels in human brains than in age-matched rodent brains. PSD-95 serves as a negative control and GAPDH serves as a loading control (Mouse n = 4; Human n = 4). **(B)** Quantification of normalized FRMPD2 chemiluminescent intensity from (A). Mean ± SEM; *P <0.05; Two tailed student’s t test. **(C)** FRMPD2 expression in rat tissues. Lysates from cortex, lung, liver, spleen, kidney, heart, and muscle were analyzed by western blot. GluA1 marks neuronal tissue; actin is a loading control. FRMPD2 is enriched in cortex. **(D)** FRMPD2 expression across mouse brain regions. Whole brain, prefrontal cortex, hippocampus, and cerebellum lysates from P90 mice were analyzed by western blot. FRMPD2 is enriched in hippocampus, similar to GluA1. Neuroligin-1, GluA1, and PSD-95 are excitatory synaptic markers; GAPDH is a loading control (n = 3 mice per region). **(E)** FRMPD2 expression in primary rat hippocampal cultures. Mixed neuron–glia or pure glial cultures were analyzed by western blot. NeuN and GFAP mark neurons and glia, respectively; GAPDH is a loading control. FRMPD2 is neuron-specific. **(F)** Developmental FRMPD2 expression in mouse brain. Lysates from P0, P5, P10, P15, P20, and P30 brains were analyzed by western blot. FRMPD2 peaks at P10, decreases at P20, and increases again at P30. PSD-95 and neuroligin-1 show increasing expression from P0 to P15. n = 4 brains per time point. **(G)** Schematic of the SLENDR system. AAV-HDR contains gRNA and GFP flanked by homology arms; AAV-EFsCas9 encodes Cas9 with nuclear localization signals and a Flag tag. The gRNA targets exon 1 of the rat FRMPD2 gene. **(H)** Genomic editing in neurons. Neurons were co-transduced with AAV-HDR and AAV-EFsCas9 at DIV9 and fixed at DIV16. PSD-95 marks excitatory synapses. Endogenous GFP-FRMPD2 co-localizes with PSD-95 in rat hippocampal neurons (arrows). **(I)** FRMPD2 enrichment in synaptosomes. Rat brain fractions were analyzed by western blot. FRMPD2 is enriched in synaptosomes, alongside synaptic proteins GluA1, PSD-95, and neuroligin-1, confirming fraction purity.

A previous RNA expression profile study showed that in humans, FRMPD2 RNA is highly expressed in brain tissue relative to other organs (Lizio 2015), while an earlier study detected FRMPD2 protein in multiple epithelial and neuronal cell lines (Stenzel 2009). However, the protein expression pattern of FRMPD2 among different organs and tissues remains less clear. To examine the FRMPD2 protein expression profile, we collected tissue from various mouse organs, including the cortex, lung, liver, spleen, kidney, heart, and muscle, and probed FRMPD2 by western blot. We found that FRMPD2 is enriched in cortical brain tissue relative to other tissue samples (**Fig. 1C**). AMPA receptor (AMPAR) subunit GluA1 was probed as a neuronal marker. We further dissected the brain regions from adult mice at postnatal day 100 (P100) and prepared lysates from the whole brain, prefrontal cortex, hippocampus, and cerebellum. Synaptic proteins, including neuroligin-1, GluA1, and PSD-95, were also probed. Western blots revealed that FRMPD2 is highly expressed in the hippocampus and prefrontal cortex, but not in the cerebellum, and its expression profile resembles that of the probed synaptic proteins, including neuroligin-1 (**Fig. 1D**). Next, because neurons and glia are two major cell types in the brain, we tested for neuronal specificity of FRMPD2. To this end, samples were prepared from either a mixed culture of neurons and glia or a culture of pure glia. Western blotting results showed that FRMPD2 was present in the lysate from a mixed culture of neurons and glia but absent from pure glial culture (**Fig. 1E**), suggesting that FRMPD2 expression is neuron-specific. We further examined the developmental profile of FRMPD2 expression in mouse brains at various time points: P0, P5, P10, P15, P20, and P30. Whole brains were lysed and subjected to western blotting. FRMPD2 expression steadily increased over time with a developmental pattern similar to the synaptic proteins, suggesting a role in synapse formation and maturation (**Fig. 1F**).

### FRMPD2 is enriched at synaptic sites in neurons

Previous study has shown that FRMPD2 localizes to the basolateral compartment within epithelial cells at the site of cell adhesion (Stenzel 2009). Because the basolateral compartment in epithelial cells is considered the counterpart of the somatodendritic compartment in neurons, we wondered whether FRMPD2 is localized at dendritic spines and implicated in neuronal adhesion at synaptic sites. In line with this, FRMPD2 contains three PDZ domains, which is a common structural hallmark for synaptic proteins such as PSD-95, Shank3 and GRIP (Dong 1997, Stenzel 2009, Harris 2001, Kim 2004).

We first wanted to examine the subcellular distribution of FRMPD2 in neurons. Due to a lack of available FRMPD2 antibodies for immunostaining of endogenous FRMPD2, we developed a CRISPR-Cas9 based gene editing tool to introduce a GFP tag specifically to the beginning of exon 1 of the FRMPD2 gene (**Fig. 1G**). Endogenous rat FRMPD2 was thus expressed as a GFP fusion protein (GFP-FRMPD2). To determine synaptic localization of FRMPD2, neurons were immunostained with the synaptic marker PSD-95. GFP-FRMPD2 showed a high degree of co-localization with PSD-95 puncta (**Fig. 1H**), suggesting a strong enrichment of FRMPD2 at the excitatory synaptic sites. To further confirm the synaptic distribution of FRMPD2 *in vivo*, we purified the synaptosome from rat cortical brain tissue and probed for FRMPD2. Western blots demonstrated an abundant presence of FRMPD2 in the synaptic fraction, together with other known synaptic proteins including PSD-95, GluA1, and neuroligin-1 (**Fig. 1I**). These results confirmed that FRMPD2 is a protein enriched in the synapse, both *in vitro* in cultured neurons and *in vivo* in the brain.

### High dosage of FRMPD2 leads to an increase in excitatory synapses

To examine the role of FRMPD2 in synaptogenesis, we transduced cultured hippocampal neurons at the time of plating with lentiviral GFP-FRMPD2 or GFP alone as a control. After 14 days, neurons were fixed and immunostained for the synaptic marker proteins synapsin and PSD95 to determine synapse density along dendrites. We found that, compared to the GFP control, neurons with FRMPD2 overexpression showed a significant increase in synaptic puncta density for both synapsin and PSD-95 (**Fig. 2A-B**). To further confirm the role of FRMPD2 in synaptogenesis, we transfected scrambled shRNA as a control and an shRNA against FRMPD2 to knockdown FRMPD2 in cultured hippocampal neurons from DIV0 to DIV14. As predicted, immunostaining for synapsin revealed a significant decrease in puncta density. Co-transfection of an shRNA targeting rat FRMPD2 together with an shRNA-resistant human GFP-FRMPD2 construct effectively rescued the knockdown phenotype (**Fig. 2C-D**). These findings strongly support a role for FRMPD2 in promoting synaptogenesis during neuronal development.

**Figure 2.**
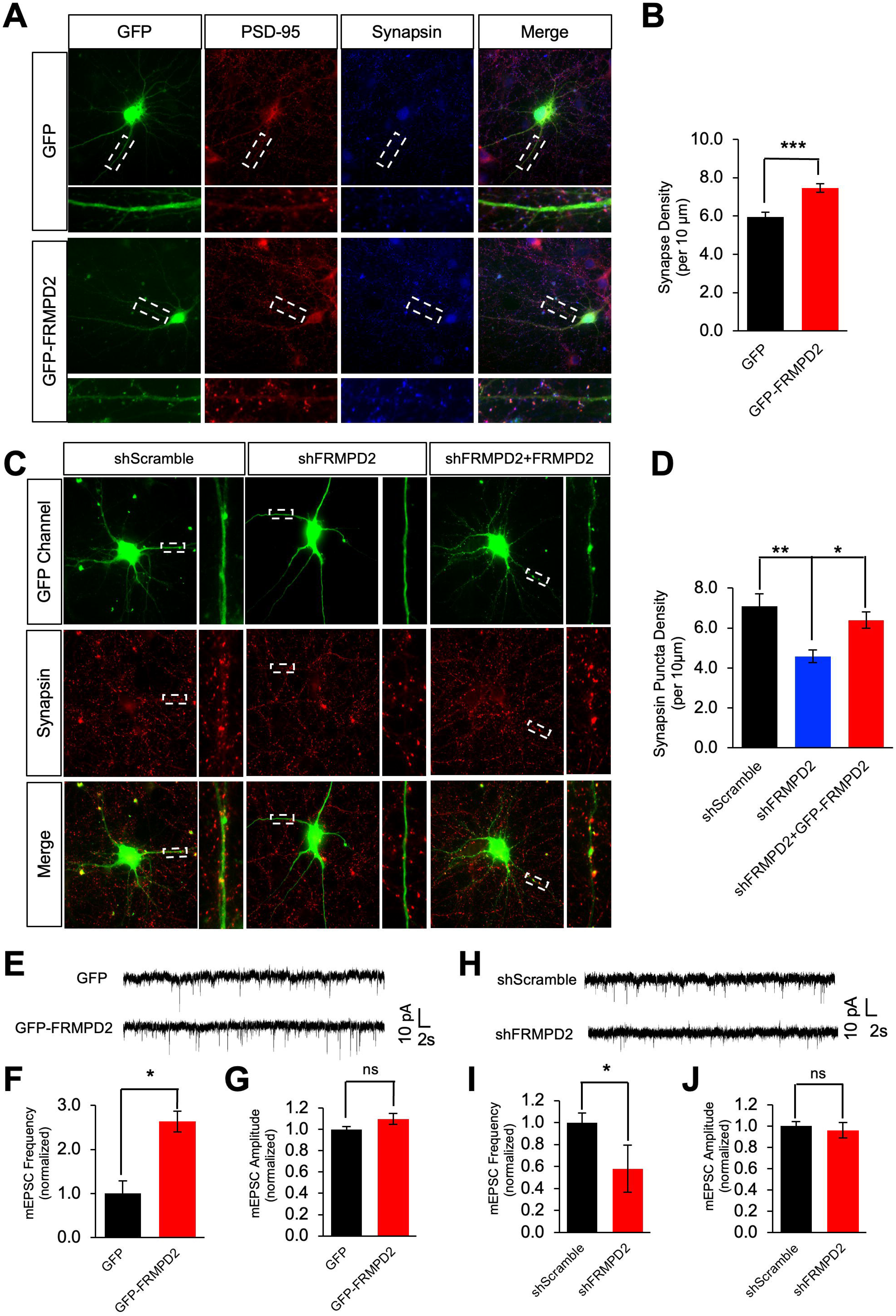
Altering FRMPD2 protein levels *in vitro* impacts synapse formation in primary rat cultured neurons. **(A)** Double immunostaining of Synapsin (presynaptic) and PSD-95 (postsynaptic) in neurons transfected with GFP or GFP-FRMPD2 at DIV11 and fixed at DIV14. Synapses were defined by co-localization of Synapsin and PSD-95. FRMPD2 overexpression increases synaptogenesis. **(B)** Quantification of co-localized PSD-95/Synapsin puncta density. n = 10 GFP & 10 GFP-FRMPD2 cells. **(C)** Immunostaining of Synapsin in neurons transfected with scrambled shRNA, shRNA targeting rat FRMPD2, or shRNA targeting rat FRMPD2 with human GFP-FRMPD2 rescue. Neurons were transfected at DIV11 and fixed at DIV14. FRMPD2 knockdown reduces Synapsin puncta density, which is rescued by human GFP-FRMPD2. **(D)** Quantification of Synapsin puncta density. shScramble (n=8 cells), shFRMPD2 (n=11 cells), and shFRMPD2+GFP-FRMPD2 (n=11 cells). **(E)** Representative miniature EPSC recording traces from neurons transfected with GFP alone or GFP together with GFP-FRMPD2. Neurons were transfected at DIV12 and recorded at DIV14. **(F-G)** Normalized mEPSC frequency (F) and amplitude (G) from neurons transfected with GFP alone or GFP together with GFP-FRMPD2. Upregulation of FRMPD2 led to an increase in mEPSC frequency without changing the amplitude. GFP (n=6 cells) and GFP-FRMPD2 (n=7 cells). **(H)** Representative mEPSC recording traces from neurons transfected with scrambled shRNA or shRNAs against FRMPD2. Neurons were transfected at DIV12 and recorded at DIV14. **(I-J)** Normalized mEPSC frequency (I) and amplitude (J) from neurons transfected with scrambled shRNA or shRNAs against FRMPD2. Downregulation of FRMPD2 led to a decrease in mEPSC frequency without changing the amplitude. shScramble (n=5 cells) and shFRMPD2 (n=5 cells). Mean ± SEM; *P <0.05, **P <0.01, ***P<0.001, ns = non-significant; Two tailed student’s t test (B,F,G,I,J) or One-way ANOVA with Tukey’s post hoc test (D).

To determine the effect of FRMPD2 on synaptic transmission, we performed patch-clamp recordings of miniature excitatory postsynaptic currents (mEPSCs). At DIV12, cultured hippocampal neurons were transfected with GFP-FRMPD2 or GFP alone and mEPSCs were recorded two days later. Compared to the GFP control, overexpression of GFP-FRMPD2 significantly increased the average mEPSC frequency while the average amplitude remained unchanged (**Fig. 2E-G**). In another set of experiments, DIV12 hippocampal neurons were transfected with either an shRNA to knockdown endogenous FRMPD2 or with a scrambled shRNA control. We found that FRMPD2 knockdown reduced mEPSC frequency while the average amplitude remained unchanged (**Fig. 2H-J**). This data supports that FRMPD2 promotes synapse formation and enhances synaptic activities.

### FRMPD2 associates with neuroligin-1 via the PDZ domain

Analysis of the FRMPD2 amino acid sequence suggests that it is an intracellular protein with no tentative transmembrane domains. Given that FRMPD2 contains multiple domains including PDZ and FERM domains, FRMPD2 is predicted to function as a scaffold protein at the synaptic domain. We hypothesize that FRMPD2 interacts with and recruits a synaptogenic membrane protein to promote synapse formation. A previous study reported that FRMPD2 interacts with the type-I PDZ binding motif at its associated proteins including δ-catenin, P0071, and ARVCF (Stenzel et al 2009). In examination of proteins involved in synapse formation, we found that neuroligin family members contain a type I PDZ binding motif (-T-T-R-V) at their C-terminus (**Fig. 3A**). Neuroligin-1 is of particular interest because it is an adhesion molecule implicated in synaptogenesis (Chubykin 2007, Futai 2013, Scheiffele 2000, Chih 2005, Craig 2007, Budreck 2013, Nam 2005). We thus speculated whether FRMPD2 regulates synapse formation through an interaction with neuroligin-1. To test this idea, we co-transfected HEK cells with GFP-FRMPD2 and HA-neuroligin-1. In cell lysates, FRMPD2 was immunoprecipitated with anti-GFP antibodies and probed with HA antibodies for neuroligin-1. The co-IP assays confirmed the association of FRMPD2 with neuroligin-1 (**Fig. 3B**). To examine whether association of neuroligin-1 with FRMPD2 affects the surface distribution of neuroligin-1, we transduced cultured cortical neurons at DIV7 with GFP-FRMPD2 or GFP control lentiviruses. 7 days later, we performed surface biotinylation to isolate the surface proteins, and samples were probed neuroligin-1 by western blot (**Fig. 3C**). Indeed, we found that the amount of cell surface neuroligin-1 was increased in GFP-FRMPD2 overexpression neurons, while the total protein level of neuroligin-1 remained unchanged compared to control GFP control neurons (**Fig. 3D**).

**Figure 3.**
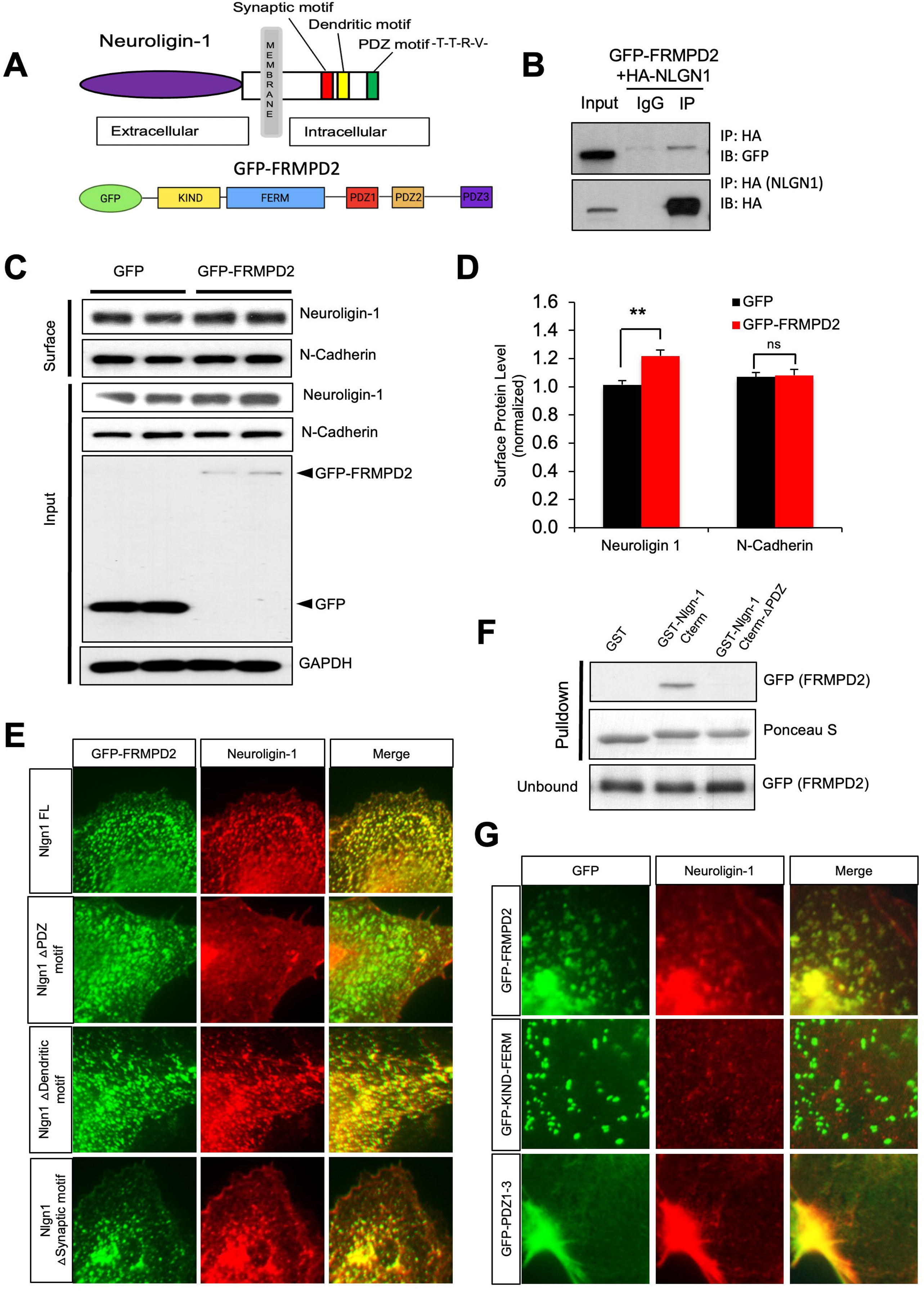
FRMPD2 associates with postsynaptic neuroligin-1 in a PDZ-PDZ binding motif-dependent manner. **(A)** Diagrams of postsynaptic neuroligin-1 and the fusion protein GFP-FRMPD2 depicting key functional domains. **(B)** Co-immunoprecipitation assay in HEK cells co-transfected with GFP-FRMPD2 and HA-Neuroligin-1 shows that Neuroligin-1 interacts with FRMPD2. **(C)** Surface biotinylation assay in primary neurons transduced with either GFP or GFP-FRMPD2 viruses. Neuroligin-1 and N-cadherin were probed from the surface fraction and total lysate. **(D)** Quantification of western blot from (C). Overexpression of GFP-FRMPD2 led to an increase in the surface expression of neuroligin-1 but not N-cadherin (n=4 independent experiments). **(E)** Fluorescent images of COS-1 cells transfected with HA-neuroligin-1 full length, HA-neuroligin-1 ΔPDZ motif, HA-neuroligin-1 Δdendritic motif or Neuroligin-1 Δsynaptic motif mutants, alone or with GFP-FRMPD2. Cells were fixed 24 h post-transfection and immunolabeled with HA antibody. Green, red, and yellow boxes indicate PDZ-binding, synaptic localization, and dendritic localization motifs, respectively. Alone, neuroligin-1 constructs show lamellipodial enrichment without puncta. Co-expression with GFP-FRMPD2 induces puncta co-localization for all constructs except the ΔPDZ mutant. **(F)** Pulldown assay showing PDZ-binding motif–dependent interaction between neuroligin-1 and FRMPD2. GST or GST fusion proteins purified from E. coli were incubated with HEK cell lysates expressing equal amounts of GFP-FRMPD2. Unbound and pulldown fractions were analyzed by western blot. **(G)** Fluorescent images of COS-1 cells transfected with GFP-FRMPD2, GFP-KIND-FERM, or GFP-PDZ1-3, alone or with HA–neuroligin-1. Cells were fixed 24 h post-transfection and immunolabeled with HA antibody; neuroligin-1 alone was included as control. GFP-FRMPD2 and GFP-KIND-FERM form intracellular puncta, whereas GFP-PDZ1-3 is diffusely distributed. GFP-KIND-FERM does not co-localize with HA–neuroligin-1, while GFP-PDZ1-3 co-localizes with HA–neuroligin-1 and is enriched in lamellipodia. Mean ± SEM; **P <0.01, ns = non-significant; Two tailed student’s t test (D).

Transmembrane proteins often use motifs at the intracellular C-terminus for interaction with intracellular partners including scaffold proteins (Calver 2001, Walther 2015). We wondered whether the interaction with FRMPD2 is mediated by the C-terminal motif of neuroligin-1. There are three known motifs in the intracellular C-terminus of neuroligin-1: the synaptic localization motif, dendritic localization motif, and a type I PDZ binding motif at the end of the terminus. To determine the motif involved in the interaction, we made truncation deletions of each individual motif. We co-transfected GFP-FRMPD2 or GFP, together with either the full length (FL) HA-neuroligin-1 or HA-neuroligin-1 with motif truncations (**Fig. 3E**). Both full-length proteins co-localized and formed discrete clusters. Similar co-clustered patterns were detected in cells co-expressing GFP-FRMPD2 and truncated neuroligin-1 with either deletion of the synaptic motif or deletion of the dendritic localization motif, indicating that these two motifs are dispensable for FRMPD2 association. In contrast, in cells expressing neuroligin-1 lacking the PDZ binding motif (-T-T-R-V), the co-patching pattern completely disappeared. These results indicate that the PDZ binding motif in neuroligin-1 mediates its interaction with FRMPD2. To further confirm the role of the PDZ binding motif in neuroligin-1, we created GST-tagged fusion proteins with full length C-terminus of neuroligin-1, or neuroligin-1 C-terminus lacking the PDZ binding motif (-T-T-R-V). An equal amount of GST, GST-NLGN1 C-term, or GST-NLGN1 C-term without the PDZ binding motif (GST-Nlgn-1 Cterm-△PDZ) were incubated with HEK293T cell lysates expressing GFP-FRMPD2. We found that only the GST-NLGN1 C-term successfully pulled down GFP-FRMPD2, indicating that the interaction is indeed mediated by the PDZ binding motif in neuroligin-1 (**Fig. 3F**).

Next, we examined the necessity of the domains in FRMPD2 for its association with neuroligin-1. In Cos-1 cells, we transfected either GFP-FRMPD2, FRMPD2 without PDZ domains (GFP-KIND-FERM), or FDMPD2 without KIND/FERM domains (GFP-PDZ1-3), together with HA-neuroligin-1. When co-expressed in Cos-1 cells, GFP-FRMPD2 and GFP-PDZ1-3, but not GFP-KIND-FERM, showed a high degree of colocalization with neuroligin-1. In addition, in co-transfected cells, GFP-KIND-FERM, but not GFP-PDZ1-3, continued to show a clustered pattern (**Fig. 3G**). These findings indicate that while the KIND-FERM domains are critical for the punctate pattern, the PDZ domains are required for FRMPD2’s interaction with neuroligin-1.

### FRMPD2 recruits neuroligin-1 to synaptic sites and promotes synapse formation

Having shown that FRMPD2 is associated with neuroligin-1, we wanted to know whether FRMPD2 regulates the distribution of neuroligin-1 in neurons. We transfected DIV11 hippocampal neurons with GFP-FRMPD2 or GFP and immunostained for endogenous neuroligin-1 on DIV 14 (**Fig. 4A**). Neuroligin-1 was distributed along the dendrites in a punctate pattern, presumably at synaptic sites. Compared to the GFP control, neuroligin-1 puncta intensity was significantly increased in neurons overexpressing GFP-FRMPD2 (**Fig. 4B**).

**Figure 4.**
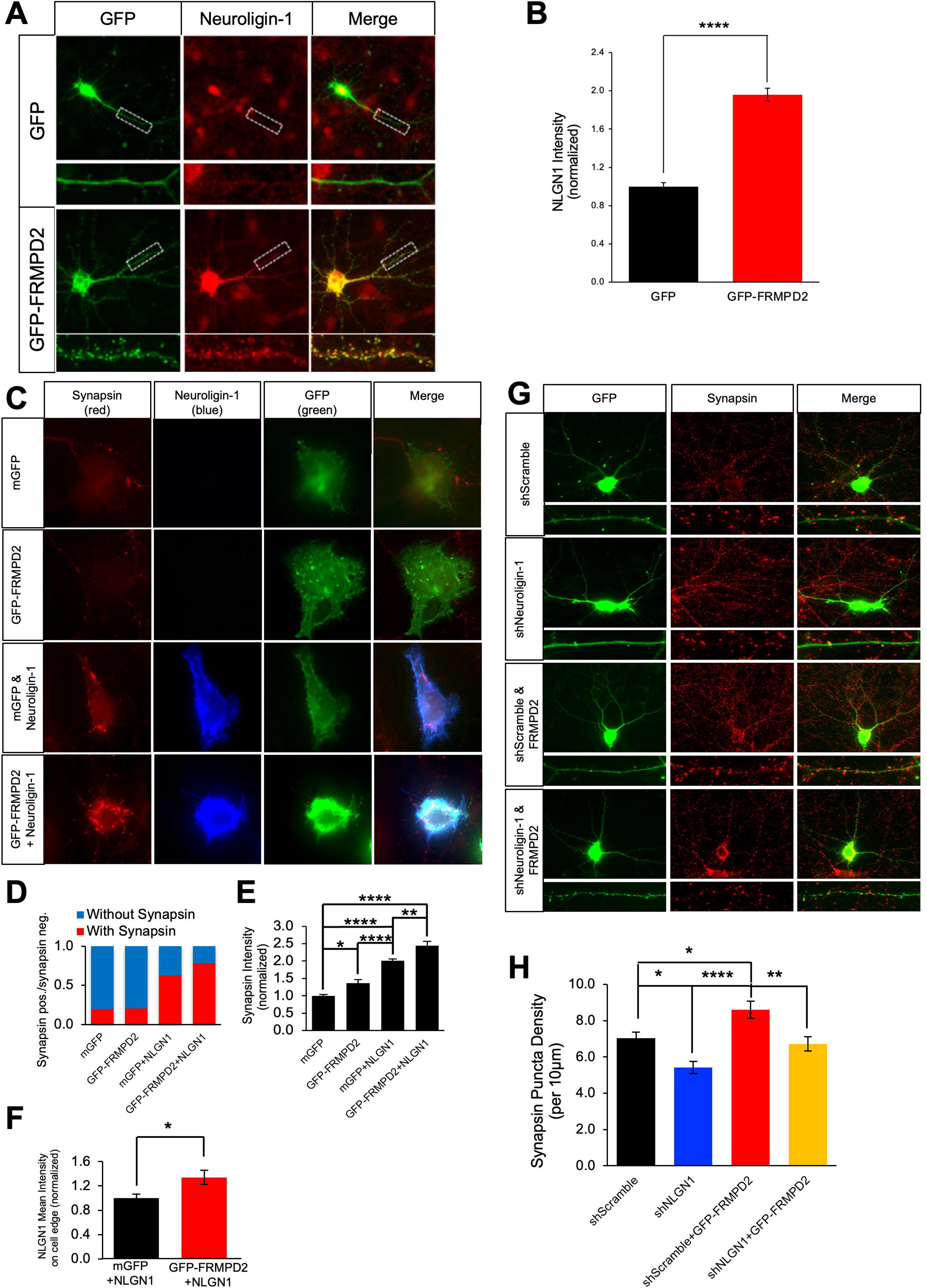
FRMPD2 requires Neuroligin-1 to functionally promote synaptogenesis at somatodendritic contact points. **(A)** Immunofluorescent staining of neuroligin-1 in primary hippocampal neurons transfected with GFP or GFP-FRMPD2. Neurons were transfected at DIV11, followed by fixation at DIV14. **(B)** Quantification of neuroligin-1 intensity from (A). Overexpression of GFP-FRMPD2 led to increased Neuroligin-1 puncta intensity (n=10 cells/condition). **(C)** Representative fluorescent images of HEK cells co-cultured with DIV10 neurons. HEK cells were transfected at DIV8 with membrane GFP, GFP-FRMPD2, membrane GFP + HA–neuroligin-1, or GFP-FRMPD2 + HA–neuroligin-1. After 24 h, cells were trypsinized and added to primary hippocampal neurons at DIV9. Co-cultures were fixed at DIV10 and immunolabeled for Synapsin and HA. **(D)** Quantification of the percentage of Synapsin-positive HEK cells. GFP or GFP-FRMPD2 alone show low Synapsin-positive cells, whereas co-expression of neuroligin-1 with mGFP increases Synapsin-positive cells. mGFP (n=30 cells), GFP-FRMPD2 (n=34 cells), mGFP+HA-neuroligin-1 (n=32 cells), and GFP-FRMPD2+HA-neuroligin-1 (n=37 cells). **(E)** Quantification of synapsin puncta intensity in Synapsin-positive HEK cells. GFP-FRMPD2 overexpression resulted in increased synapsin puncta intensity within the area of HEK cells compared to mGFP alone. mGFP+HA-neuroligin-1 overexpression led to increased Synapsin puncta intensity compared to GFP-FRMPD2. mGFP (n=10 cells), GFP-FRMPD2 (n=10 cells), mGFP+HA-Neuroligin-1 (n=8 cells), and GFP-FRMPD2+HA-Neuroligin-1 (n=9 cells). **(F)** Quantification of mean neuroligin-1 intensity on the edge area of HEK cells transfected with either mGFP+HA-neuroligin-1 or GFP-FRMPD2+HA-Neuroligin-1. The presence of GFP-FRMPD2 increased the mean intensity of HA-neuroligin-1 on the cell edge. mGFP+HA-neuroligin-1 (n=28 cells) and GFP-FRMPD2+HA-neuroligin-1 (n=33 cells). **(G)** Synapsin immunostaining in neurons transfected from DIV 11 to 14 with scrambled shRNA alone, neuroligin-1 shRNA alone, scrambled shRNA + GFP-FRMPD2, or neuroligin-1 shRNA + GFP-FRMPD2. **(H)** Quantification of synapsin puncta density from transfected neurons in (G). Neuroligin-1 knockdown reduced Synapsin puncta density, while GFP-FRMPD2 overexpression led to its increase. The FRMPD2-induced increase in Synapsin was abolished by neuroligin-1 knockdown. Mean ± SEM; *P <0.05, **P <0.01, ****P<0.0001, ns = non-significant; Two tailed student’s t test (B,F) or One-way ANOVA with Tukey’s post hoc test (E,H).

As a synaptic adhesion molecule, neuroligin-1 is known to induce synaptic formation via its interaction with the presynaptic protein Neurexin (Craig 2007, Nam 2004, Levinson 2005, Scheiffele 2000). Based on our findings showing that FRMPD2 interacts with neuroligin-1 and recruits neuroligin-1 to the cell surface, we hypothesized that FRMPD2 may play a role in pre- and postsynaptic adhesion and synapse formation. To test this hypothesis, we performed synaptogenic assays on co-cultures of neurons and heterologous cells. Briefly, we transfected HEK cells with GFP-FRMPD2 or GFP, together with or without HA-neuroligin-1. One day after transfection, HEK cells were collected and placed on top of DIV 9 cultured hippocampal neurons grown on coverslips. One day later, cells were fixed and immunostained with antibodies against synapsin and HA (for HA-neuroligin-1). Consistent with previous studies (Levinson 2005, Scheiffele 2000), we found that expression of neuroligin-1 in HEK cells led to an increase in synapsin-positive HEK cells (**Fig. 4C-D**). Interestingly, co-expression of FRMPD2 and neuroligin-1 resulted in a significant further increase in the number of synapsin-positive HEK cells (**Fig. 4D**). In contrast, HEK cells expressing only GFP or GFP-FRMPD2 had a minimal basal level of synapsin signals, indicating that FRMPD2 alone was unable to attract presynaptic terminals to HEK cells. In line with changes in synapsin puncta, neuroligin-1 overexpression also increased synapsin intensity, which was further increased in cells co-expressing GFP-FRMPD2 (**Fig. 4E**). At the peripheral edge area of HEK cells, neuroligin-1 intensity was increased by 50% due to the presence of GFP-FRMPD2 compared to neuroligin-1 alone (**Fig. 4F**). These findings indicate that FRMPD2 and neuroligin-1 synergistically attract presynaptic terminals, presumably functioning to facilitate synapse formation.

We showed that FRMPD2 interacts and co-distributes with neuroligin-1, and expression of either protein led to an increase in synapse numbers. The positive regulation of synaptogenesis could result from independent pathways downstream of FRMPD2 or neuroligin-1, or these two proteins may function sequentially in the same cascade leading to synapse formation. To determine the functional dependency of these molecules, we transfected DIV11 neurons with an shRNA against neuroligin-1 or scrambled shRNA as control, with or without co-transfection of FRMPD2. We found that knockdown of neuroligin-1 reduced synapsin puncta density compared to the scramble shRNA control. More importantly, knockdown of neuroligin-1 blocked the increase in synapsin puncta density induced by FRMPD2 (**Fig. 4G-H**). These results suggest that the FRMPD2 effect on synaptogenesis is mediated by neuroligin-1.

To evaluate the involvement of specific PDZ domains in FRMPD2, we transfected neurons at DIV11 with GFP-FRMPD2, GFP-KIND-FERM, GFP-PDZ1-3, GFP-FRMPD2 ΔPDZ1 (i.e., GFP-FRMPD2 lacking PDZ1), GFP-FRMPD2 ΔPDZ2 and GFP-FRMPD2 ΔPDZ3 and stained neuroligin-1. We found that GFP-FRMPD2, GFP-FRMPD2 ΔPDZ1, GFP-FRMPD2 ΔPDZ2 or GFP-FRMPD2 ΔPDZ3 were able to increase neuroligin-1 puncta density and intensity, whereas the truncated GFP-KIND-FERM showed no effect (**Fig. 5A-C**). These data indicate that all three PDZ domains, but not the KIND/FERM domains, are required for the effect of FRMPD2 on neuroligin-1 synaptic recruitment.

**Figure 5.**
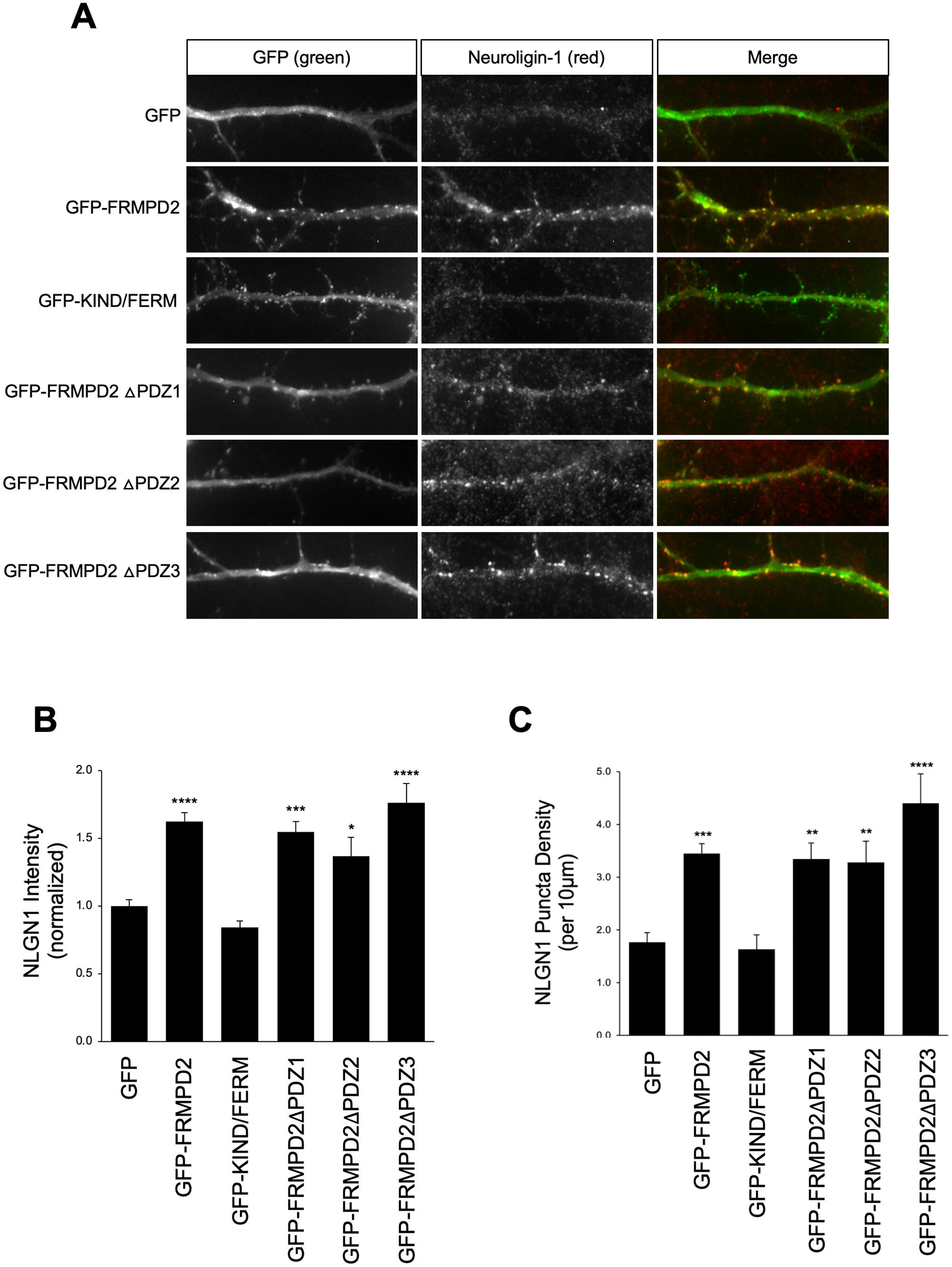
The PDZ and FERM domains in FRMPD2 are required to enrich Neuroligin-1 at synaptic sites. **(A)** Representative images of dendrites from neurons transfected with GFP alone as control, or together with GFP-FRMPD2, GFP-KIND-FERM, GFP-PDZ1-3, GFPFRMPD2 ΔPDZ1, GFP-FRMPD2 ΔPDZ2 or GFP-FRMPD2 ΔPDZ3, respectively. Cells were immunostained for Neuroligin-1. **(B)** Quantification of neuroligin-1 puncta intensity from (A) revealed that the FERM domain and PDZ domains were sufficient to induce enrichment of Neuroligin-1. GFP (n=13 cells), GFP-FRMPD2 (n=17 cells), GFP-KINDFERM (n=14 cells), GFP-PDZ1-3 (n=11 cells), GFP-FRMPD2 ΔPDZ1 (n=12 cells), GFP-FRMPD2 ΔPDZ2 (n=11 cells), and GFP-FRMPD2 ΔPDZ3 (n=11 cells). **(C)** Quantification of neuroligin-1 puncta density from (A) revealed that with the FERM domain, any two PDZ domains were sufficient to induce enrichment of Neuroligin-1. GFP (n=13 cells), GFP-FRMPD2 (n=17 cells), GFP-KINDFERM (n=14 cells), GFP-PDZ1-3 (n=11 cells), GFP-FRMPD2 ΔPDZ1 (n=12 cells), GFPFRMPD2 ΔPDZ2 (n=11 cells), and GFP-FRMPD2 ΔPDZ3 (n=11 cells). Mean ± SEM; *P <0.05, **P <0.01, ***P <0.001, ****P <0.0001; One-way ANOVA with Dunnett’s post hoc test (B,C).

### FRMPD2 promotes spine formation and morphological maturation

Spines are categorized into three types: mushroom, stubby, and thin spines. Mushroom and stubby spines are often considered as mature spines, while thin spines are considered as an immature type. To test whether FRMPD2 contributes to spine maturation, DIV11 hippocampal neurons were transfected with either GFP-FRMPD2 or GFP together with membrane-GFP to better visualize the spines (**Fig. 6A**). At DIV 14, neurons were imaged, and spine density and subtypes were analyzed. Compared to control conditions, overexpression of GFP-FRMPD2 led to an increase in overall spine density, with no change in spine head width or spine length (**Fig. 6B-F**). Of the spine subtypes, the mushroom and thin spines, as well as filopodia, were increased, while the stubby spine density remained unchanged. These data suggest that a high dosage of FRMPD2 promotes spine formation and spine morphological maturation. To further determine the role of FRMPD2 on spinogenesis, we utilized shRNA-mediated knockdown of endogenous FRMPD2. DIV11 cultured rat hippocampal neurons were transfected with GFP and either a mixture of shRNAs against FRMPD2 or scrambled shRNA as a control. GFP-positive cells were used for spine density analysis. Compared to the control, FRMPD2 knockdown led to a significant reduction in spine density (**Fig. 6G-H**).

**Figure 6.**
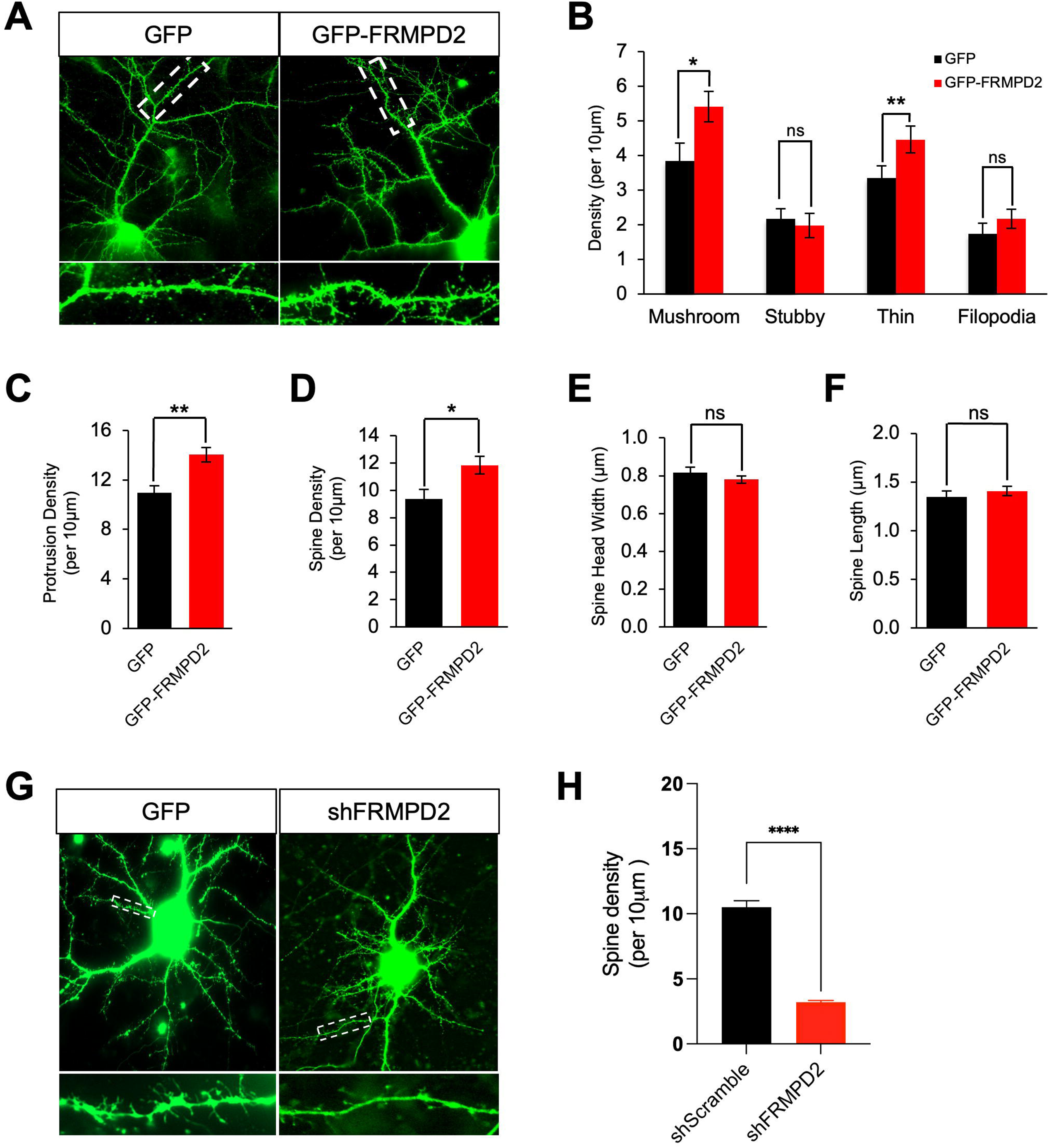
FRMPD2 promotes dendritic spine maturation in primary rat cultured neurons. **(A)** Representative fluorescent images of neurons transfected with mGFP alone or mGFP together with GFP-FRMPD2. Representative enlarged dendritic segments are shown below the neuron. Neurons were transfected at DIV11, followed by fixation at DIV14. **(B)** Quantification of mGFP-labeled protrusion density. Protrusions were classified into mushroom spines, stubby spines, thin spines and filopodia. Overexpression of FRMPD2 increased mushroom and thin spine density, while stubby spine and filopodia density remained unchanged. **(C-F)** Quantification of protrusion density (C), spine density (D), spine head width (E), and spine length (F). Overexpression of FRMPD2 resulted in elevated protrusion density and spine density without changing spine head width and spine length. GFP (n=11 cells) and GFP-FRMPD2 (n=12 cells). **(G-H)** Representative fluorescent images (G) and spine density quantification (H) of neurons transfected from DIV11 to 14 with mGFP alone or mGFP + FRMPD2 shRNA. FRMPD2 knockdown reduced spine density. n = 10 GFP cells and 10 shFRMPD2 + GFP cells. Mean ± SEM; *P <0.05, **P< 0.01, ****P< 0.0001, ns = non-significant; Two-tailed student’s t test (B-F, H).

### Association of FERM domain with F-actin leads to FRMPD2 synaptic localization

FRMPD2 is a protein with multiple functional domains: a KIND domain at the N-terminal, a FERM domain, and three PDZ domains at the C-terminal. Our results found synaptic enrichment of FRMPD2, but the molecular elements responsible for its subcellular distribution remain unknown. We thus wanted to determine the role of each FRMPD2 domain for its synaptic localization.

The FERM domain has been shown to interact with the actin cytoskeleton in Myosin1 (Gotesman 2010) and in Talin (Lee 2004). Because actin is highly enriched in the spine (Hotulainen 2010, Landis 1983), we wondered whether F-actin could serve as an anchor to dock FRMPD2 at the synaptic sites. To test whether FRMPD2 associates with actin, we co-transfected HEK293T cells with either GFP-FRMPD2 or GFP control together with mCherry-actin for 2 days, and the cell lysates were used for co-IP assays (**Fig. 7A**). Western blotting revealed that a high level of actin exists in the immunoprecipitants of FRMPD2, indicating an association between FRMPD2 and actin. To map the domain of FRMPD2 responsible for this interaction, HEK cells were transfected with truncated FRMPD2, including KIND-FERM, PDZ1-3, KIND-only, FERM-only and FERM deletion, respectively, together with mCherry-actin or GFP as a control. Consistent with our hypothesis, co-IP showed that mCherry-actin was co-immunoprecipitated with GFP-KIND-FERM or FERM-only, but not with the other truncations. These data indicate that the FERM domain in FRMPD2 may mediate the association with actin (**Fig. 7B**).

**Figure 7.**
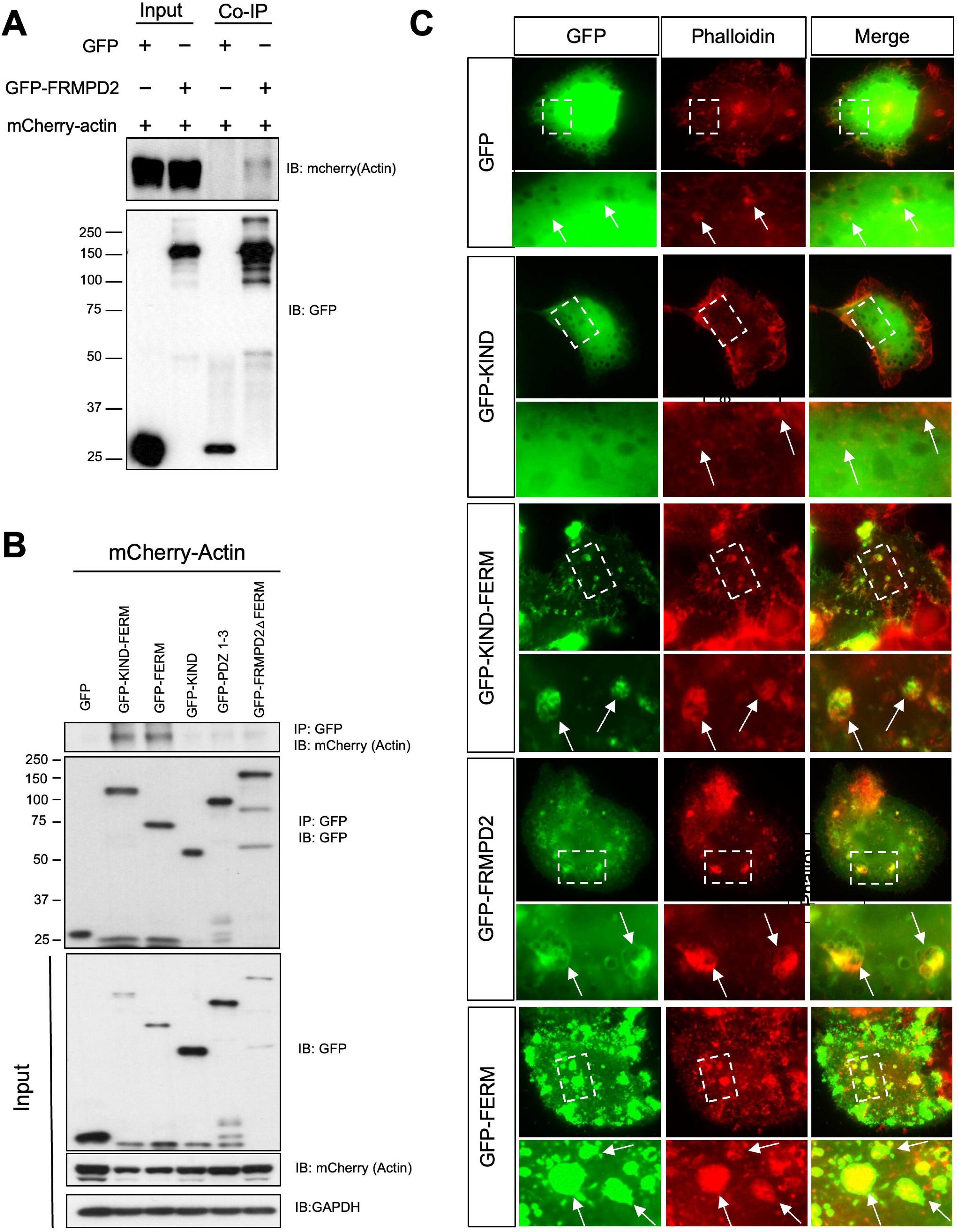
FRMPD2 interacts with F-Actin through the FERM domain to facilitate accurate localization. **(A)** Co-immunoprecipitation assay between GFP-FRMPD2 and mCherry-actin in HEK cells. Western blot analysis confirmed the association between FRMPD2 and actin. **(B)** Co-immunoprecipitation assay between mCherry-actin and GFP or GFP fused with various FRMPD2 truncations, including GFP-KIND-FERM, GFPFERM, GFP-KIND, GFP-PDZ1-3 and GFP-FRMPD2 ΔFERM in HEK cells. Western blot revealed an interaction between the FERM domain and mCherry-actin. **(C)** Phalloidin-labeled F-actin fluorescent images of HEK cells transfected for 24 hours with GFP, GFP-FRMPD2, GFP-KIND-FERM, GFP-KIND or GFP-FERM, respectively. GFP-FRMPD2, GFP-KIND-FERM and GFP-FERM formed co-patching with F-actin. GFP and GFP-KIND showed ubiquitous presence without co-patching formation.

To further confirm the interaction of the FERM domain with actin, we transfected HEK cells with GFP, GFP-FRMPD2, GFP-KIND, GFP-KIND-FERM, and GFP-FERM, respectively, and labeled F-actin with fluorophore-conjugated phalloidin (**Fig. 7C**). Our results showed that GFP-FRMPD2, GFP-KIND-FERM and GFP-FERM colocalized with F-actin at the peripheral region of the cell and co-patched with F-actin within the cells. Interestingly, expression of GFP-FERM led to a reorganization of the F-actin pattern, forming co-patching plaques within HEK cells containing both GFP-FERM and distorted F-actin. These results indicate the necessity of the FERM domain for the FRMPD2 interaction with F-actin, and suggest a reorganization of F-actin patterns following complexing with FRMPD2 in cells.

To examine the role of F-actin on FRMPD2 in neurons, rat hippocampal neurons were transfected with GFP-FRMPD2 or GFP at DIV11, and at DIV13, neurons were treated with either DMSO as a control or Latrunculin A to disassemble the synaptic F-actin network. Fixation and labeling of F-actin with fluorescent phalloidin at DIV14 showed that Latrunculin A treatment completely disrupted F-actin filaments within neurons when compared to DMSO control (**Fig. 8A**). Accordingly, both F-actin puncta density and intensity decreased significantly. Consistent with changes in F-actin, GFP-FRMPD2 also showed a reduction in puncta size, intensity, and density (**Fig. 8B-D**). These data support the idea that the enrichment of F-actin in the spine leads to the recruitment of FRMPD2 to the synaptic compartment.

**Figure 8.**
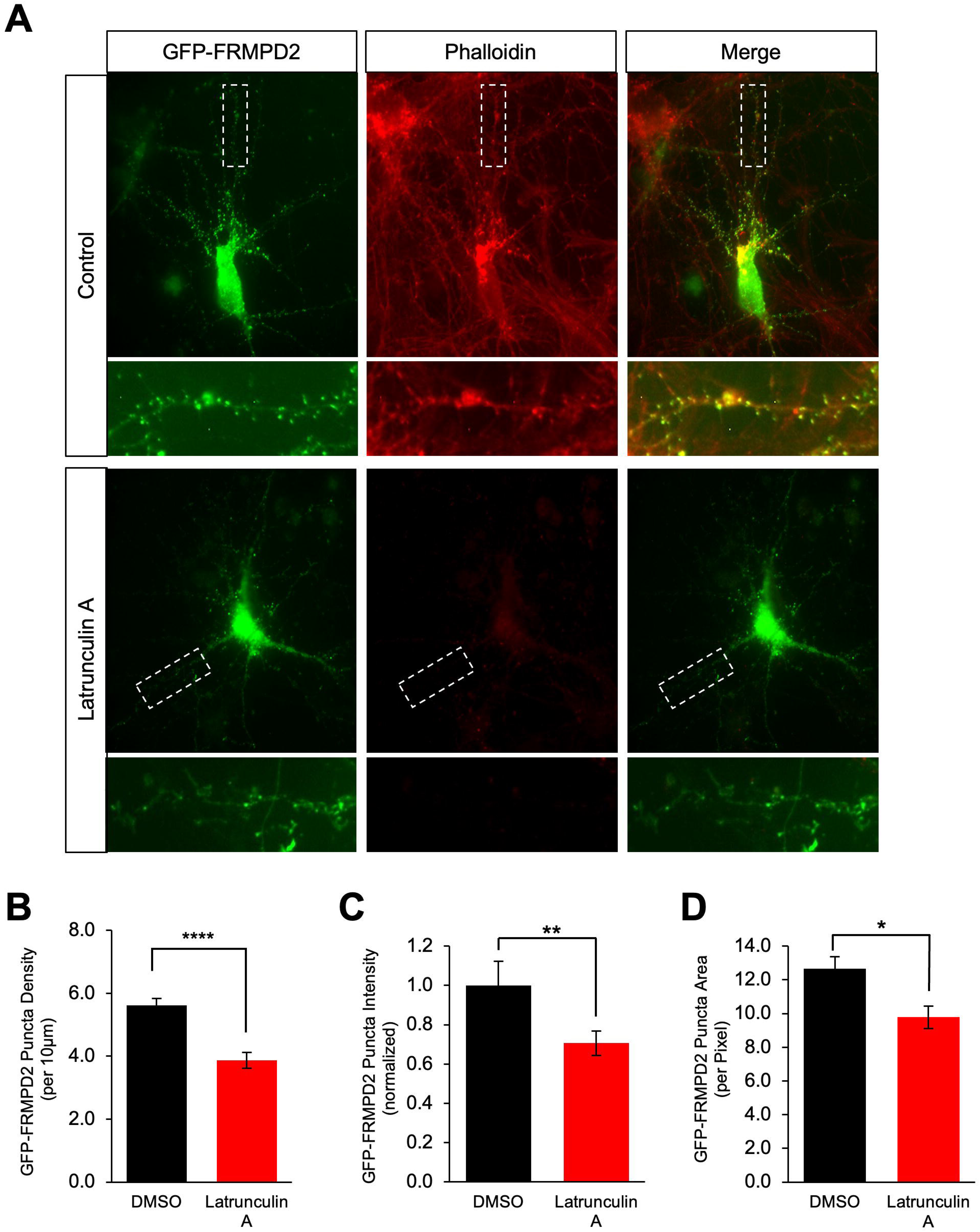
The FRMPD2 domains FERM and PDZ are required for accurate spine-specific and synaptic localization through interaction with the F-actin network. **(A)** Fluorescent images of primary hippocampal neurons transfected with GFP-FRMPD2 at DIV11 and treated with either DMSO or 5μM Latrunculin A at DIV13. Neurons were fixed at DIV14 and subjected to subsequent F-actin labeling by fluorophore-conjugated phalloidin. **(B-D)** Quantification of GFP-FRMPD2 puncta density (B), puncta intensity (C), and puncta area (D) revealed that depolymerization of the F-actin network by Latrunculin A treatment reduced GFP-FRMPD2 puncta density, puncta intensity, and puncta area. DMSO (n=14 cells) and Latrunculin A (n=14 cells). Mean ± SEM; *P <0.05, **P <0.01, ****P <0.0001. Two tailed student’s t test (B-D).

### FRMPD2 overexpression leads to delayed neuronal migration *in vivo* in the mouse brain

Our results demonstrated a higher level of FRMPD2 expression in human brains than in rodents, and a high dosage of FRMPD2 via overexpression promotes synapse formation in primary rat neurons. We wondered whether FRMPD2 affects other neurodevelopmental processes in the brain *in vivo*. To this end, we performed *In utero electroporation* (IUE) in embryonic mouse brains, where a small group of neurons were electroporated with a mixture of GFP-FRMPD2 and RFP or RFP only as a control. IUE was performed on Embryonic Day 14.5 (E14.5) to label neurons that would eventually migrate to cortical layer II/III. Following birth, mouse brains were collected at either P0 or P15 and fixed for slicing and imaging. We found that at P0, neurons with FRMPD2 overexpression had a higher fraction of cells distributed in the SVZ and IZ regions, and a lower fraction localized to the cortical plate, compared to the control condition (**Fig. 9A-B**). At P15, control neurons had mostly migrated to layer II/III and the majority of FRMPD2 overexpressing neurons expectedly resided in the upper cortical plate. However, a small portion of FRMPD2 overexpressing neurons were found in the lower cortical plate (**Fig. 9C-D**). These data suggest that FRMPD2 hyperexpression in neurons results in delayed neuronal migration during brain development *in vivo*.

**Figure 9.**
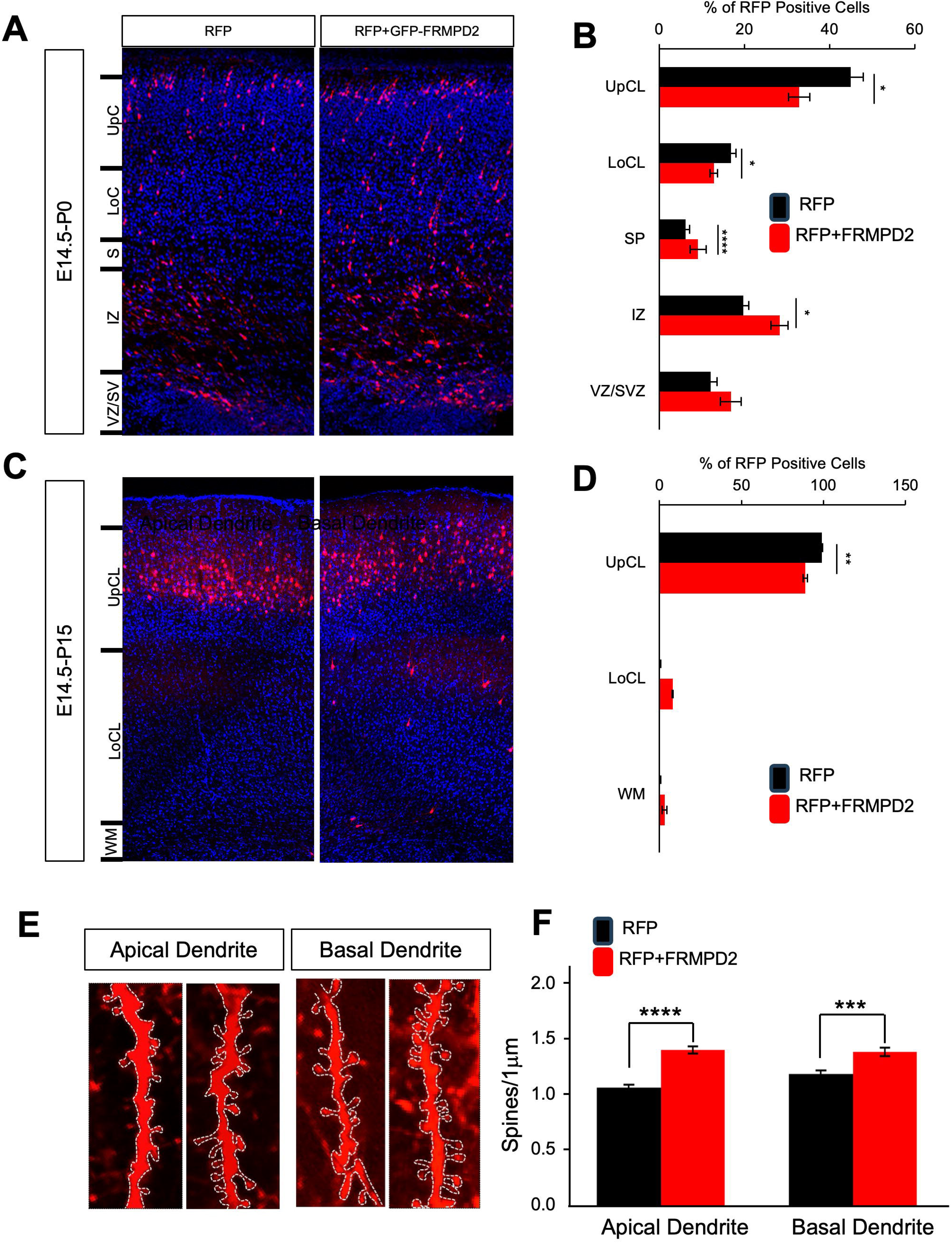
FRMPD2 overexpression leads to delayed neuron migration and increased spine density during mouse brain development. **(A)** Representative coronal brain slice images from P0 mouse brains following in utero electroporation (IUE) at E14.5 with RFP alone as a control or RFP together with GFP-FRMPD2. **(B)** Quantification of the percentage of RFP-positive cells in the upper cortical layer (UpCL), lower cortical layer (LoCL), subplate (SP), intermediate zone (IZ), and ventricular/subventricular zones (VZ/SVZ). The percentage of FRMPD2-overexpressing neurons was reduced in both the UpCL and LoCL, but higher in the IZ. n=4 Control brains and 4 FRMPD2 brains. **(C)** Representative coronal brain slice images from P15 mouse brains following IUE at E14.5 with RFP alone as a control or RFP together with GFP-FRMPD2. **(D)** Quantification of the percentage of RFP-positive cells in the UpCL, LoCL, and white matter (WM) revealed that the percentage of FRMPD2-overexpressing neurons was reduced in the UpCL and a fraction was localized to the LoCL. n=3 Control brains and 3 FRMPD2 brains. **(E)** Representative dendrites from the Layer II/III neurons of P15 mouse brains following IUE at E14.5. **(F)** Quantification of basal and apical spine density from (F) revealed that overexpression of FRMPD2 increased both basal and apical spine density. Apical: Control (n=36 cells) and GFP-FRMPD2 (n=37 cells); Basal: Control (n=30 cells) and GFP-FRMPD2 (n=26 cells). Mean ± SEM; *P <0.05, **P <0.01, ***P <0.001, ****P <0.0001. Two-tailed student’s test (B,D,F).

We also analyzed the effect of FRMPD2 overexpression on spine formation in the apical and basal dendrites. We found that as compared to control, FRMPD2 overexpression resulted in increased spine density on both apical and basal dendrites (**Fig. 9E-F**). These data further support a role for FRMPD2 on promoting spine and synapse formation as observed *in vitro*.

### High dosage of FRMPD2 in the brain results in enhanced spatial memory in mice

Synaptogenesis and neuronal connectivity play a critical role in brain functions including cognition (Huo 2022, Gilbert 2020, Gatto 2010, Betancur 2009, Irwin 2005). Given our findings demonstrating the significant synaptogenic effects of FRMPD2, we wanted to know whether elevated FRMPD2 expression in the brain affects animal behavior. To this end, we adopted an adeno-associated virus (AAV) to globally overexpress HA-FRMPD2, or GFP as a control, in mouse brains. In P0 mouse pups, high-titer viruses were intraventricularly injected into the lateral ventricles. 2 months later, the injected mice of both sexes were subject to Barnes Maze test to determine the effects of FRMPD2 on spatial learning and memory. We first measured the primary latency to reach the escape hole and the primary hole distance during acquisitions. We found that both measurements were not significantly different (**Fig. 10A-B**), suggesting that the learning capacity remained unchanged in FRMPD2 overexpression mice. During the 24-hour post-test and 5-day post-test, the target hole was sealed, and animals were placed on the maze to explore and locate the target hole. We found that FRMPD2 overexpression mice spent more time in the target quadrant and more time at the proximity of the target hole than controls (**Fig. 10C-H**), indicating improved memory retention in these animals. These data suggest that a high dosage of FRMPD2 in the brain contributes to enhanced memory capability in rodents.

**Figure 10.**
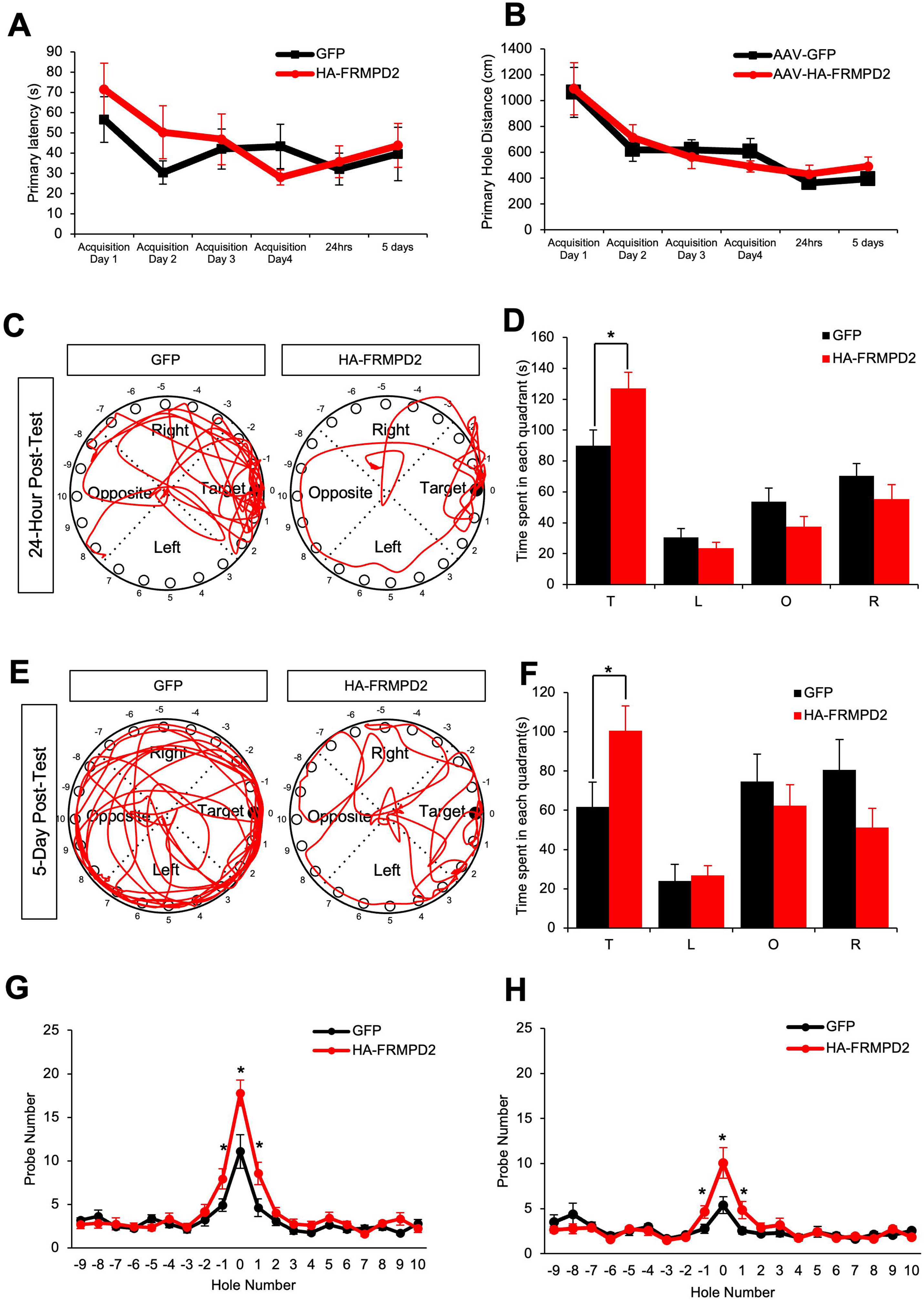
AAV-facilitated overexpression of FRMPD2 in the developing mouse brain improves memory retention in adults. **(A-B)** Quantification of the primary latency to reach the target hole (A) and the primary hole distance (B) in the Barnes maze assay revealed no difference between mice overexpressing GFP (n=13) or HA-FRMPD2 (n=14) during training and testing. **(C)** Representative trajectories of mice overexpressing GFP or HA-FRMPD2 during the 24-hour post-test. **(D)** Quantification of time spent in each quadrant during the 24-hour post-test revealed that HA-FRMPD2 overexpressing mice spent significantly more time in the target quadrant than GFP controls. **(E)** Representative trajectories of mice overexpressing GFP or HA-FRMPD2 during the 5-day post-test. **(F)** Quantification of time spent in each quadrant during the 5-day post-test revealed that HA-FRMPD2 overexpressing mice spent significantly more time in the target quadrant than GFP controls. **(G-H)** Quantification of nose probe number during the 24-hour (G) and 5-day (H) post-test revealed that HA-FRMPD2 overexpressing mice had a higher average probe number on target holes “-1”, “0”, and “1” than that of GFP controls. Mean ± SEM; *P <0.05; Two tailed student’s t test (A,B,D,F,G,H).

## DISCUSSION

As a multidomain scaffold protein, the neurobiological role of FRMPD2 remains less clear. We found that FRMPD2 is preferentially expressed in the brain and enriched at synapses. Its synaptic localization appears to be anchored by a FERM domain–mediated association with F-actin. Through its PDZ domains, FRMPD2 interacts with and recruits neuroligin-1 to postsynaptic sites, thereby promoting synaptogenesis. Thus, FRMPD2 functions as a pivotal coordinator linking the dendritic spine actin network to the key synaptic adhesion molecule neuroligin-1. Consistently, enhanced neuronal connectivity is associated with improved cognitive performance in spatial memory.

The FERM domain of FRMPD2 interacts with F-actin and promotes its reorganization, while F-actin depolymerization by latrunculin A disrupts FRMPD2 localization and reduces FRMPD2 puncta density. Conversely, the interaction between neuroligin-1 and FRMPD2 is PDZ domain-and PDZ binding motif-dependent, and all three PDZ domains in FRMPD2 are equally important for this association. This interaction leads to a higher level of neuroligin-1 at the surface and at synaptic sites. Co-culture experiments show that while neuroglin-1 alone facilitates synapse formation, neuroligin-1 and FRMPD2 work synergistically to maximize the effect. These findings suggest that FRMPD2 functions as a scaffold protein that bridges the actin cytoskeleton to synaptic adhesion molecules at the postsynaptic domain, and consequently promotes synaptogenesis. In addition, FRMPD2 has been shown to interact with NMDARs at excitatory synapses through its second PDZ domain (Lu 2019). Together with NMDAR signaling at the post-synapse through PSD-95 (Sheng 2001), these suggest that FRMPD2 acts as a hub molecule in the formation of a functional protein network at the synapse.

Neuroligin-1 is a key postsynaptic adhesion molecule required for synaptogenesis and functional synaptic signaling. Dysregulation of neuroligin proteins has been implicated in several neurological disorders (Lee et al., 2023). Mutations in neuroligin-1 have been reported in individuals with autism spectrum disorder (ASD) (Rajabi et al., 2024; Nakanishi et al., 2017). At the synapse, neuroligin-1 is tightly regulated by multiple mechanisms to maintain proteostasis and proper function. Notably, neuroligin-1 mRNA undergoes local translation at synapses in response to NMDAR-driven activity, a process mediated by the fragile X mental retardation protein (FMRP) (Chmielewska et al., 2019). Loss of FMRP results in increased surface expression of neuroligin-1 and enhanced synaptic transmission. Our results demonstrate that FRMPD2 recruits neuroligin-1 to synaptic sites through its PDZ domain, which in turn promotes synaptogenesis. It has also been reported that the PDZ2 domain of FRMPD2 interacts with NMDARs, with a higher preference for the synaptic GluN2A subunit than the extra synaptic GluN2B subunit, while the full length FRMPD2 protein is required to coprecipitate with the GluN1/N2A functional complex (Lu 2019). Furthermore, NMDAR-mediated activity-driven proteolytic cleavage of neuroligin-1 has been previously established as a mechanism in regulating excitatory synapse plasticity (Suzuki 2012). Our study sheds light on a scaffold protein which acts as a central regulator of neuroligin-1 proteostasis at the postsynaptic site. Thus, FRMPD2 could be a key molecular target for designing therapeutics aiming to control synapse formation and neural circuit connectivity.

Reflecting a similar line of thought, a recent therapeutic approach to virally deliver a functional copy of SHANK3 to brain cells to rescue SHANK3 haploinsufficiency has progressed to Phase I/II clinical trials (Betancur 2013, Costales 2015, National Library of Medicine, 2024). Recent studies in developing engineered synaptic adhesion molecules that can promote precise synaptogenesis with high specificity and are able to organize synaptic proteins into functional synapses (Hale 2022, Rabinowitch 2024) have been of recent importance in this area of research. The modular use of extracellular and intracellular domains of synaptic adhesion molecules can enable directed and controlled synaptogenesis as a potential intervention for neurodevelopmental disorders. Furthermore, a synthetic extracellular adhesion molecule, CPTX, designed with domains interacting with cerebellin-1 and neuronal pentraxin-1, exhibits strong potency as a synthetic synaptic organizer at excitatory synapses (Suzuki et al 2020). Further structural classification of the FRMPD2 protein and its complexes with other adhesion molecules can provide greater depth into precise chemical motifs to target and develop therapeutic strategies.

Our *in vivo* FRMPD2 overexpression experiments show increased spine densities in layer II/III pyramidal neurons compared to control cells. Moreover, the primary dendrite number in FRMPD2 overexpression neurons was also increased, suggesting a role of FRMPD2 in dendrite morphogenesis during neuronal development. Spine and synapse density, as well as the overall size of the dendritic arbor, are markedly higher in humans than in primates and other mammals (Mohan 2015). Studies have reported that human pyramidal neurons receive more synaptic inputs than other primates and rodents (DeFelipe 2011, Elston 2001). Human cortical evolution is also signified by, in comparison to rodents, an increase in total synapse number per neuron in all cortical layers except layer IV, and therefore enhanced connectivity during brain development and expansion (DeFelipe 2011). Given the higher level of FRMPD2 expression in human brains, FRMPD2 may contribute to the unique features in human brain development. In line with this, our study shows that overexpression of FRMPD2 in primary rat neurons leads to a significant increase in spine and synapse formation.

Interestingly, FRMPD2 overexpressing mice at both P0 and P15 showed a delay in neuronal migration. Comparative anatomical studies reveal that humans demonstrate a protracted period of development consisting of a slower migration speed (Rao 2001) and multiple protein families interact to bring about carefully controlled cortical layer formation through radial glia-guided migration (Nascimento 2023, Alderman 2024, Geschwind 2013, Rakic 1988, Rakic 2003). Further studies have found that migration mutations that cause severe disorders in the human brain cause only subtle phenotypes in rodents, emphasizing the importance of migration and lamination in proper human brain development (Buschbaum 2019, Gleeson 2000). On a functional level, numerous cell-adhesion molecules and cytoskeleton interacting genes are implicated in neural migration disorders (Buschbaum 2019). Given FRMPD2’s interaction with cell adhesion molecules and actin, it may become a potential neural migration target for further study, yet future studies are needed to elucidate the molecular mechanism underlying this effect. Recent development of cerebral organoid strategies and controllable gene expression are tools that can enable these studies.

Comparative genomic sequencing studies show that FRMPD2 is one of the human-specific multi-copy genes compared with other primates (Sudmant 2010). Regions with segmental duplications (SD) are likely to undergo non-allelic homologous recombination (NHAR), which could introduce microduplications, misalignments, and deletion mutations at the SD genomic loci, which give rise to copy number variation (CNV) disorders (Stankiewicz 2012) and gene dosage-related phenotypes. Indeed, both copies of FRMPD2 in the human genome reside in 10q11.21-q11.23, a region reported to have duplication or deletion associated with neurological deficits including ASD, intellectual disability (ID), and developmental delays (DD) in human patients (Tritto 2021, Fu 2021, Stankiewicz 2012). Genetic studies revealed that the segment harboring FRMPD2 and PTPN20 duplicated about 2.3 million years ago during evolution and left 2 copies of FRMPD2 in the human genome (Sudmant 2010). Recent genetic and experimental studies on the FRMPD2 gene have revealed that FRMPD2 copies derive their novel Transcriptional Start Site (TSS) from an internal promoter, as the native TSS is lost during segmental duplication (Dougherty 2018), leading to the formation of 5’ truncations (Dennis 2017). One such truncated copy, FRMPD2B, has been described as a paralog with neural expression and involvement in brain development (Soto 2025). Our study found that the protein levels of full length FRMPD2 are higher in human brain tissues than in mouse brain, suggesting that the elevated expression of FRMPD2 in the human brain may not be purely due to extra gene copies, rather, it might result from an upregulation in FRMPD2 transcription or suppression of FRMPD2 protein turnover. Further proteomic studies involving the full length and truncated forms of FRMPD2 as well as the distinct functional isoforms produced by these genes will be required to better understand evolutionary control of proteostasis and genetic duplications. Hence, FRMPD2 may be a prime candidate to understand the evolution of higher complexity observed in the human brain.

### Limitations of the study

This work was mostly conducted in primary cultured neurons. Further work in animal models will be instrumental to study FRMPD2 under physiological conditions. The *in vivo* experiments performed in this project were through IUE or viral elevation of gene dosage; thus, an endogenous knock-in line overexpressing full-length FRMPD2 will be an essential tool that, if developed, can provide better insight into neurodevelopmental mechanisms in wild-type rodents and their humanized transgenic counterparts. While this study revealed the importance of the FRMPD2 association with neuroligin-1, further characterization of FRMPD2’s interaction with other adhesion molecules may provide greater depth into its regulatory function in synapse formation and maturation. Moreover, our study focuses on the function of FRMPD2 function at excitatory synapses, but a more comprehensive study involving inhibitory synapses would shed light on neuronal regulation of excitatory/inhibitory synapse specificity.

## Author contributions

H.Y.M. conceived the research; H.Y.M. and Y.H. designed most of the experiments; Y.H., A.P., K.M. and J.G. performed the experiments; Y.H. analyzed most of the data, A.P. analyzed the electro-recording data, A.P. and K.M. analyzed some immunostaining data, Y.H., A.P. and H.Y.M. prepared the figures; and H.Y.M., A.P. Y.H. and K.M. wrote the paper. All authors read and approved the final manuscript.

## ACKNOWLEDGEMENTS

We would like to thank the Man Lab members for their insightful input. We thank Dr. Kai S. Erdmann (Ruhr-University Bochum, Germany) for providing FRMPD2 plasmid. We thank Viola Monovich for imaging assistance and Natalie Kochova for transfection assistance. This work was supported by R01 MH079407, R01 MH130600, R21 MH133014, and R21 MH134174.

## MATERIALS & METHODS

### Neuronal Culture Preparation and Transfection

Primary rat (Sprague Dawley) cortical and hippocampal neuron cultures were prepared from embryonic day 18 (E18) rats of either sex. Dissected cortices and hippocampi were digested with papain (0.5mg/ml in EBSS) and plated on poly-L-lysine coated coverslips or petri dishes. Neurons were plated in plating medium (MEM containing 10% fetal bovine serum, 5% horse serum, 31mg L-cysteine, and 1% penicillin/streptomycin and L-glutamine mixture (1% P/S/G); Invitrogen). After 24 hours, the plating medium was replaced by a feeding medium (Neurobasal medium supplemented with 1% HS, 2% B-27, and 1% P/S/G). Neurons were then maintained in the feeding medium and regularly fed before being used. One week after plating, 5-fluorodeoxyuridine (FDU, 5μM) was added to the media to inhibit glial growth. All cells were maintained in a humidified incubator containing 5%(v/v) CO2. Transfections were performed at different timepoints in vitro and incubated for 3 hours before media change. All transfections were performed with Lipofectamine 2000 (Invitrogen) according to the manufacturer’s instructions.

### HEK and Cos-1 Cell Cultures

Human embryonic kidney (HEK) 293A cells were cultured in Dulbecco’s Modified Eagle Medium (Gibco) supplemented with 10% heat-inactivated fetal bovine serum and 1% penicillin/streptomycin and passaged at 100% confluency twice a week. Transfections were performed at approximately 50-70% confluency using Lipofectamine 2000. One day following transfection, cells were fixed and immunostained or lysed for biochemical analysis.

Cos-1 cells were cultured in Dulbecco’s Modified Eagle Medium (Gibco) supplemented with 10% heat-inactivated fetal bovine serum and 1% P/S and passaged at 100% confluency twice a week. When the Cos-1 cell culture reached 100% confluency, the culture medium was removed, and cell cultures were washed once with pre-warmed sterile phosphate-buffered saline (PBS). Cell cultures were trypsinized with 1ml trypsin-EDTA for 3 minutes at 37°C to allow the full digestion. To neutralize the reaction, 1ml of culture medium was applied, and the mixture was triturated up and down gently by a Pasteur pipette to fully disassociate individual cells. The desired number of cells were transferred into a new dish with cell culture medium for further experiments or maintenance.

### HEK cell and Neuron Co-culture

HEK293T cells cultured in a 12-well plate were transfected separately, and cultured hippocampal neurons at DIV8 (days in vitro) were prepared in parallel. Overexpression of target proteins in HEK293T cells was allowed for 24 hours. HEK293T cells were then briefly trypsinized for 30 seconds at 37°C and triturated to separate the monolayer culture into individual cells. The cell suspension was diluted 10-fold and added to the cultured hippocampal neurons at the age of DIV9. After 24 hours of co-culture within conditioned feeding medium, both neurons and HEK293T cells were subject to fixation and immunostaining.

### CRISPR-Cas9-based Genomic Knock-in of GFP-FRMPD2

To minimize off-target effects, the guide RNA and Cas9 were packaged into separate AAV vectors, ensuring that genomic double-strand breaks were generated only in neurons co-transduced with both vectors. To enable recombination, a GFP coding sequence flanked by ∼1 kb homology arms corresponding to regions of FRMPD2 surrounding the desired knock-in site was combined with a guide RNA sequence in the vector to facilitate homology-directed insertion. Cultured rat hippocampal neurons (DIV10) were transduced with high-titer viruses encoding the Cas9 gene and guide RNA sequence and left to incubate for 7 days prior to fixation. Neurons cultured on coverslips at DIV9 were placed in a 12-well plate with 500μl of conditioned feeding medium. 10μl of AAV2-U6-gRNA-GFP-FRMPD2 virus was added together with 5μl of AAV2-NLS-Cas9-NLS-Flag into each well and incubated at 37°C for 24 hours. On DIV10, another 500μl of conditioned feeding medium was added to the well and neurons were cultured at 37°C for one week. On DIV16, neurons were fixed and subject to immunostaining. Neurons receiving both viral constructs exhibited a GFP knock-in at the start of exon 1 of rat FRMPD2.

### Synaptosome Preparation

Cortical tissue dissected from adult rat brains was minced and homogenized in an ice-cold solution (0.32M sucrose, 1mM NaHCO3, 1mM MgCl2, 0.5mM CaCl2). Lysates were transferred to 15ml conical tubes and further solubilized with the same buffer for 30 minutes at 4°C. Samples were centrifuged at 1,400 x g for 10 minutes and the supernatant (S1) was transferred to a new tube and centrifuged at 13,800 x g for 10 min. The pellet (P2) containing the synaptosome was resuspended in radioimmunoprecipitation (RIPA) lysis buffer (50mM Tris-HCl pH 7.4, 150mM NaCl, 1% NP40, 1% SDOC, 0.1% SDS). Protein concentration was measured using the BCA protein determination kit (Thermo Scientific), and both the lysate and P2 fraction were diluted to the same protein concentration with RIPA lysis buffer. 2X Laemmli sample buffer was then added followed by sample denaturation at 95°C for 10 minutes.

### Neuron and Brain Sample Preparation

Cultured neurons or HEK293T cells were lysed and collected in 2X Laemmli sample buffer (100mM Tris-HCl pH 6.8, 4% Sodium dodecyl sulfate (SDS), 0.2% bromophenol blue, 20% glycerol, 5% (v/v) β-mercaptoethanol). Samples were then incubated at 95°C for 10 minutes prior to western blotting or storage at −20°C for long-term use.

Brain tissues from humans or rodents were dissected on ice and immersed in pre-chilled RIPA buffer (50 mM Tris-HCl pH 7.4, 150 mM NaCl, 1% NP-40, 1% sodium deoxycholate (SDOC), 1% sodium dodecyl sulfate (SDS)) supplemented with a protease inhibitor cocktail tablet (Roche). For 1g tissue, 3 ml of pre-chilled RIPA buffer (1% SDS) was used. Then, tissues were homogenized mechanically with a Teflon pestle (Thermo-Fisher Scientific) and sonicated for 10 cycles of a 10-second sonication and a 10-second rest followed by rotation at 4°C for 30 minutes for further protein solubilization. Tissue lysates were then centrifuged for 25 minutes at 12000 rpm. The supernatant was transferred carefully into a new pre-chilled tube on ice and subjected to BCA assay to determine the protein concentration. All samples were then mixed with an equal volume of 2X Laemmli sample buffer. Protein levels were then normalized to 2 - 5μg/μl by adding 1X Laemmli sample buffer.

### SDS-PAGE gel electrophoresis

Protein amounts and modifications were assessed by SDS-polyacrylamide gel electrophoresis (PAGE) based on molecular weight and charge. The MiniProtein III Electrophoresis Cell (Biorad) system was used for electrophoresis according to the manufacturer’s instructions. The monomer stock solution of acrylamide/bisacrylamide (30% polyacrylamide) was purchased from Biorad and stored at 4°C. 10%(w/v) Ammonium persulfate (APS) solution was freshly prepared before casting the gel. A 4X separating gel buffer (1.5M Tris-HCl, pH 8.8) and a 4X stacking gel buffer (0.5M Tris-HCl, pH 6.8) were made in the lab and stored at room temperature. 1X running buffer was made from 10X running buffer [Tris base (0.25 M), Glycine (1.92 M), SDS (1% w/v)] and used for electrophoresis. 5X Laemmli sample buffer (250mM Tris-HCl pH 6.8, 10% SDS, 0.5% bromophenol blue, 50% glycerol, 12.5%(v/v) β-mercaptoethanol) was made and stored at −20°C in 1ml aliquots. For all experiments, 8% SDS-PAGE gels were adopted for the analysis of protein amount or modifications unless otherwise specified.

### Immunoprecipitation Assays

Brain tissues and cultured cells were lysed in 1X RIPA buffer supplemented with a protease inhibitor cocktail tablet (Roche) and a phosphatase inhibitor tablet (Roche) to avoid protein degradation and removal of phosphoryl groups. Stringent RIPA buffer (1% SDS) was adopted for immunoprecipitation (IP) assays to minimize the remaining contaminant proteins on the resin. Mild RIPA buffer (0.1% SDS) was applied for co-immunoprecipitation (Co-IP) assays to test protein interactions. Processed lysates were incubated with desired antibodies and pre-washed protein A/G agarose resin (Santa Cruz Biotechnology) for 3 hours or overnight at 4°C. Agarose resin was then washed 3 times with RIPA (1%) for IP assays or 5 times with NP-40 buffer (25 mM Tris-HCl pH7.4, 150mM NaCl, 1mM EDTA, 1% NP-40) for Co-IP assays. The agarose resin was incubated with 2X Laemmli sample buffer at 95°C for 10 minutes prior to western blotting.

### GST Pulldown Assays

GST and GST-tagged protein plasmids were transformed into *E.coli* BL21 cells and cultured as described previously. When the OD600 reached 0.6-0.8, Isopropyl β-D-1thiogalactopyranoside (IPTG, 0.25 mM) was added to induce protein expression overnight at 16°C with agitation. Cell pellets were then collected by centrifugation at 12000rpm for 5 minutes and resuspended in lysis buffer (20mM Tris-HCl pH 7.5, 1% Triton X-100, 1μg/ml DNAse, 1mg/ml lysozyme) freshly supplemented with phenylmethylsulfonyl fluoride (PMSF, 1mM) and protease inhibitor cocktail tablet (Roche). The cell suspension was then incubated at 30°C for 30 minutes and followed by sonication for 10 cycles of a 20-second sonication and a 20-second rest on ice. The lysate was subjected to centrifugation at 12000rpm at 4°C for 25 minutes. The supernatant was transferred and mixed with pre-equilibrated glutathione-Sepharose 4B (Sigma-Aldrich) beads in the column. The mixture was incubated at 4°C for 2 hours and the beads were washed once each with wash buffer A (20mM Tris-HCl pH7.5, 1% Triton X-100), wash buffer B (20mM Tris-HCl pH 7.5, 0.5% Triton X-100), and wash buffer C (20mM Tris-HCl pH 7.5, 0.1% Triton X-100). The beads were then resuspended in PBS and placed on ice for a pulldown assay or with 30% glycerol and frozen at −80°C for long-term storage. GFP-FRMPD2 was overexpressed in HEK293T cells by transfection for 2 days. Cells were then collected in RIPA buffer (0.1% SDS) supplemented with protease inhibitor cocktail tablets and subjected to sonication 5 times for 10 seconds each. The lysate was solubilized by rotation at 4°C for 30 minutes prior to centrifugation at 12000rpm at 4°C. The supernatant containing GFP-FRMPD2 was transferred into a new pre-chilled tube and placed on ice. For the pull-down assay, the same volume of lysate containing GFP-FRMPD2 was mixed with a comparable amount of GST or GST-tagged proteins, respectively. The bead amount was equalized by adding an adequate volume of fresh beads before incubation for accurate comparison. The beads were incubated with HEK293T cell lysate overnight followed by 4 washes in NP-40 buffer. 2X Laemmli sample buffer was added to the beads and boiled at 95°C for 10 minutes prior to western blotting.

### Immunostaining

Neurons or HEK cells cultured on coverslips were washed once with PBS and fixed at room temperature for 10 minutes with fixation solution (4% sucrose, 4% paraformaldehyde in PBS). The cells were then permeabilized in PBS with 0.3% SDS at room temperature for another 10 minutes.

Cells were then blocked in PBS with 10% goat serum (Gibco) at room temperature for 1 hour. Primary antibodies were diluted in PBS with 2.5% goat serum (1:200) and added to the cells for overnight incubation at 4°C. The cells were washed three times with PBS, followed by a 1-hour incubation at room temperature in Alexa Fluor-conjugated secondary antibodies diluted in PBS (1:500). The nuclei were stained with DAPI (1:10000 in PBS) and the cells were then washed twice with PBS and mounted onto slides with Prolong Gold antifade mountant (Thermo).

### Microscopy and Image Collection

After mounting, immunostained coverslips were kept in the dark for 4 hours prior to imaging. Images were collected with an inverted fluorescence microscope at a 63x with an oil-immersion objective (Zeiss Axiovert 200M). Exposure time for the fluorescence signal was first set automatically by the software, then adjusted manually so that the signals were within the full dynamic range. Either the glow scale look-up table or the histogram was used to monitor the saturation level. When analyzed using Image J software, an adequate threshold was applied to all images to select GluA1 puncta for quantitative measurements.

### Spine Analysis

Transfected neurons were chosen from coverslips at 63x in Zeiss Axiovert 200M microscope. Images were contrasted to allow the full visible range of spines. The Dendritic Spine density plug-in on ImageJ was used to identify and categorize spines. Three to five dendrites were manually traced and defined per neuron. A selection threshold of 70% was used and erroneous or missing spine selection was corrected manually. Mushroom spines are those with <2μm in length, >0.5 μm in width and a clear neck structure; stubby spines are with <2μm in length, >0.5 μm in width but without a clear neck structure, whereas thin spines are <2μm in length, <0.5 μm in width with a clear, long neck. Lastly, filopodia are defined as >2μm in length, <0.5 μm in width and without a clear neck structure.

### Puncta Density Analysis

Immunostained neurons were chosen from coverslips at 63x using a Zeiss Axiovert 200M microscope. Images were analyzed with equivalent thresholds across a set of neurons in an experiment across all conditions. ImageJ was used to manually define dendrites of choice in a selected channel.

### mEPSCs whole-cell patch-clamp recordings

Hippocampal neurons were transfected at DIV10 with either enhanced GFP (EGFP) and pcDNA3.1, or EGFP together with either FRMPD2 or shFRMPD2 to overexpress or knockdown FRMPD2, respectively. Total cDNA amounts were balanced by adding pcDNA3.1 in control transfections. 48 hours following transfection, a coverslip of neurons was transferred to a recording chamber with the extracellular solution containing 140mM NaCl, 3mM KCl, 1.5mM MgCl2, 2.5mM CaCl2, 11 glucoses and 10 HEPES (pH 7.4) which was supplemented with tetrodotoxin (TTX) (1μM) to block action potentials, 2-amino-5-phosphonopentanoate (APV) (50μM) to block NMDA receptors and bicuculline (20μM) to block GABAA receptor-mediated inhibitory synaptic currents. Whole-cell voltage-clamp recordings were made with patch pipettes filled with an intracellular solution containing 100mM Cs-methanesulfonate, 10mM CsCl, 10mM HEPES, 0.2mM EGTA, 4mM Mg-ATP, 0.3mM Na-GTP, 5mM QX-314 and 10mM Na-phosphocreatine (pH 7.4) with the membrane potential clamped at −70 mV. Recordings were performed in turn on the same day and coverslips were from the same batch of cells that were transfected at the same time. mEPSCs were recorded with an Axopatch 200B amplifier and displayed and recorded digitally on a computer for subsequent off-line analysis with Clampfit (Molecular Devices).

### In Utero Electroporation (IUE)

In utero electroporation (IUE) was performed on timed-pregnant CD-1 mice (Charles River Laboratories) at embryonic day 14.5 (E14.5). Animals were anesthetized via intraperitoneal (IP) injection of ketamine/xylazine solution (Ketamine: 20mg/ml, xylazine: 2mg/ml). For a 10g animal, 100μl of ketamine/xylazine solution was used. The fur on the abdomen was shaved and the operating field on the abdomen was sterilized with 10% povidone-iodine (Purdue Products L.P.) and an alcohol pad soaked with 70% ethanol. Then the uterine horns of the pregnant animals were exposed by midline laparotomy. 2μl of plasmid DNA with 0.1% Fast Green (Sigma-Aldrich) at a concentration of 2 μg/μl was injected into one lateral ventricle through the uterine wall and sac using a pulled glass micropipette. The anode of a tweezertrode (Harvard Apparatus) was placed over the dorsal telencephalon above the uterine muscle and pulses of 35V with 50ms duration separated by a 950ms interval were applied with a BTX ECM830 pulse generator (Harvard Apparatus). Following electroporation, the uterine horns were gently returned to the abdomen, the cavity was filled with warm PBS, and the incisions were closed with absorbable sutures, followed by a closure of the skin with silk sutures. The animal was returned to its cage and monitored closely during the recovery period. After birth, brains were collected at either P0 or P15. For brain collection, pups were anesthetized in a CO2 chamber for 5 minutes. P0 pups were decapitated, and brains were fixed within the skull. For P15 mice, animals were transcardially perfused with ice-cold 4% paraformaldehyde in PBS, the brains were removed and fixed in 4% paraformaldehyde in PBS at 4°C overnight. After fixation, brains were immersed in 30% sucrose in PBS at 4°C for 2 days. Brains were transferred to the slicing mold and embedded in OCT medium (Tissue-Tek). P0 brains were sliced at 35μm, while P15 brains were sliced at 60μm with a Leica CM 1850 cryostat (Leica Biosystems) at −20°C. All procedures were reviewed and approved by the Boston University Institutional Animal Care and Use Committee (IACUC).

### Barnes Maze Spatial Memory Test

The circular maze, 122cm in diameter, was made from a 2cm thick, white plastic board. Twenty holes 5cm in diameter were equidistantly drilled around the perimeter of the maze, with a distance of 2.5 cm from the edge. The maze was mounted on a pedestal allowing rotation around its center and lifted 76 cm above the ground. The escape cage, made from a black plastic box with a ramp for easy access, was placed beneath the escape hole under the maze. Four overhead fluorescent lights and a noisy buzzer served as the aversive stimuli during the test. The surface of the maze and escape cage were thoroughly wiped with 70% ethanol and 2% Virkon between trials for disinfection, and to avoid any olfactory cues affecting the animal behavior. During the test, the whole maze was isolated from the surroundings using a white curtain with four distinct pictures as visual cues for spatial directions. The mice were habituated to the maze on day 1. Each time, one mouse was placed in the center of the maze, covered by an opaque cardboard chamber for 10 seconds, then manually and carefully guided to the escape hole. Each mouse was given 5 minutes to enter the escape hole on its own, and if not, the mouse was encouraged to escape into the hole by gently pulling its tail in the direction away from the hole. Once entering the hole, all fluorescent lights and the buzzer were immediately turned off, and the animal was allowed to rest in the escape cage for 2 minutes. From day 2 to day 4, all animals went through a similar procedure, excluding the step of guidance toward the location of the escape hole, and were video recorded for learning assessment. In addition, animals were allowed to rest in the escape cage for 1 minute during acquisition days. Memory retention was tested 24 hours after and 6 days after all acquisition sessions. Before the memory retention tests, the escape hole was covered, and the maze was rotated to eliminate possible odor cues without changing the position of the visual cues. During the memory retention tests, animals were allowed to freely explore the maze for 5 minutes and were again recorded. The animal movement was traced using MouseActivityMaster MATLAB software for accuracy along with manual measurement of the primary latency and duration of stays in each quadrant.

### Statistical Analysis

Intensity signals in western blotting images were analyzed, measured, and quantified by ImageJ. The same area within each lane was selected and measured in terms of grey intensity. Background was subtracted before comparison. Independent experiments were performed and thus all of the statistical analyses were based on results derived from these experiments.

For all immunostaining images, at least three segments of secondary dendrites in length from 60μm to 100μm from different dendrites were analyzed to represent one neuron. The same threshold was used when measuring a batch of neuron images.

The MouseActivityMaster package in MATLAB was used for behavioral experiment tracing. BORIS was used for manual behavior scoring.

The GraphPad PRISM statistical analysis software was used for two sample t-Tests and analysis of variance (ANOVA) tests. Outliers were removed using robust outlier test (ROUT) with a false detection rate (FDR) value of 1%.

